# Molecular basis for Gβγ-mediated activation of phosphoinositide 3-kinase γ

**DOI:** 10.1101/2023.05.04.539492

**Authors:** Chun-Liang Chen, Ramizah Syahirah, Sandeep K. Ravala, Yu-Chen Yen, Thomas Klose, Qing Deng, John J. G. Tesmer

## Abstract

The conversion of PIP2 to PIP3 by phosphoinositide 3-kinase γ (PI3Kγ) is a critical step in neutrophil chemotaxis and is essential for metastasis in many types of cancer. PI3Kγ is activated via directed interaction with Gβγ heterodimers released from cell-surface G protein-coupled receptors (GPCRs) responding to extracellular signals. To resolve how Gβγ activates PI3Kγ, we determined cryo-EM reconstructions of PI3Kγ–Gβγ complexes in the presence of various substrates/analogs, revealing two distinct Gβγ binding sites, one on the p110γ helical domain and one on the C-terminal domain of the p101 subunit. Comparison of these complexes with structures of PI3Kγ alone demonstrates conformational changes in the kinase domain upon Gβγ binding similar to those induced by Ras·GTP. Assays of variants perturbing the two Gβγ binding sites and interdomain contacts that change upon Gβγ binding suggest that Gβγ not only recruits the enzyme to membranes but also allosterically controls activity via both sites. Studies in a zebrafish model examining neutrophil migration are consistent with these results. These findings set the stage for future detailed investigation of Gβγ-mediated activation mechanisms in this enzyme family and will aid in developing drugs selective for PI3Kγ.

---

Phosphoinositide (PI) 3-kinases (PI3Ks) are lipid kinases that phosphorylate PI lipids at the 3’-OH group to produce PI(3)P, PI(3,4)P2, and PI(3,4,5)P3. Although most are coupled to the activation of receptor tyrosine kinases (RTKs)^1^, some are also regulated by G protein-coupled receptors (GPCRs). One such isoform is PI3Kγ ^2^ whose activity was first noted in activated neutrophils after formyl peptide stimulation (f-Met-Leu-Phe)^3^. The enzyme is primarily expressed in immune cells, where it converts PI(4,5)P2 (PIP2) to PI(3,4,5)P3 (PIP3) and triggers membrane recruitment of PIP3-dependent effectors such as P-Rex1/2 and AKT. These enzyems initiate downstream signaling pathways involved in actin remodeling, cell growth, proliferation, and cell migration^4,5^. Like other Class I PI3Ks, PI3Kγ consists of a catalytic subunit (p110γ) and a regulatory subunit (p84 or p101). Lack of p84 expression in neutrophils results in reduced production of reactive oxide species (ROS), whereas lack of p101 significantly impairs chemoattraction and migration of leukocytes^6^. Aberrant expression and activation of PI3Kγ has been linked to the progression of rheumatoid arthritis, atherosclerosis, cardiovascular diseases, lupus, and pulmonary fibrosis^7^. Increased expression of PI3Kγ has also been observed in pancreatic and prostate cancer^8,9^.

PI3Kγ is stimulated by heterotrimeric Gβγ heterodimers and Ras·GTP, which are released after activation of GPCRs and RTKs^10^, respectively. However, beyond membrane recruitment, the molecular mechanisms underlying activation by Gβγ are unclear. The p110γ catalytic subunit consists of an adaptor binding domain (ABD), a Ras-binding domain (RBD), a C2 domain, a helical domain (HD), and a kinase domain (**Fig. 1A**). At the outset of our study, the structure of p101 was unknown, but was recently shown shown to consist of three domains which we refer to as the N-terminal domain (NTD), central domain (CD), and C-terminal domain (CTD) (**Fig. 1A**)^11^. Early studies strongly suggested that Gβγ binds to both p101 and p110γ subunits ^12,13^, with the p101 domain interaction seeming to be more important for membrane recruitment of PI3Kγ ^13^. The two binding sites were further localized using hydrogen-deuterium exchange mass spectrometry (HDX-MS), which pointed to the p110γ HD and the p101 CTD being the two domains responsible for Gβγ interaction^11,14^. However, ambiguities remain because changes in HDX-MS signals do not always register direct binding nor can they differentiate between direct binding events or consequent allosteric events. To clarify the regulation of PI3Kγ by Gβγ, we determined cryo-EM reconstructions of PI3Kγ in complex with Gβγ, confirming the existence of two discrete Gβγ binding sites. For comparison, we also etermined a 3.0 Å structure of heterodimeric PI3Kγ in its native conformation. Gβγ binding to the p110γ subunit drives conformational changes that serve to open the kinase active site. Based on homology, we predict that PI3Kβ, another isoform regulated by Gβγ^15^, will also engage and be regulated by Gβγ at the analogous site. Gβγ binding to the p101 CTD in our system seems dependent on Gβγ binding to p110γ and promotes further opening of the kinase active site, consistent with the two binding sites being allosterically coupled. Because these structural changes occur in the absence of membrane, it supports the idea that Gβγ is important for not only membrane recruitment but also conformational regulation independent of the membrane. Functional studies conducted *in vitro* and in a zebrafish animal model support theses conclusions..

**Figure 1.**
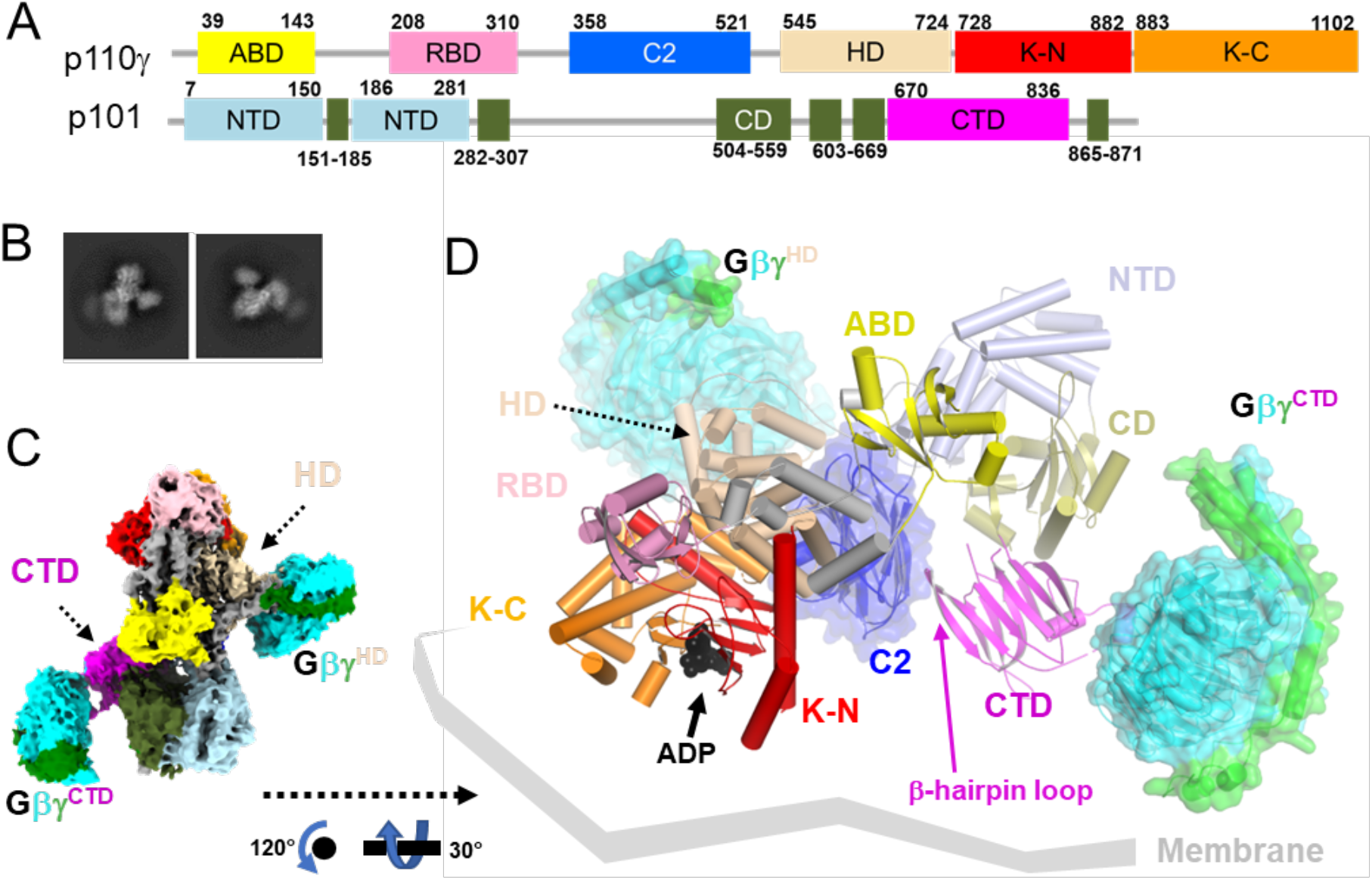
Cryo-EM reconstruction of the PI3Kγ·ADP–Gβγ^HD^–Gβγ^CTD^ complex. **(A)** Domain structure of the PI3Kγ heterodimer. The p110γ subunit is composed of an adaptor binding domain (ABD, yellow), Ras-binding domain (RBD, pink), C2 domain (blue), helical domain (HD, wheat), kinase domain N-lobe (K-N, red), and kinase domain C-lobe (K-C, orange). The p101 subunit is composed of an N-terminal domain (ND, lightblue), a central domain (CD, deepolive), and C-terminal domain (CTD, magenta). **(B)** Representative 2D class averages, roughly corresponding to the views in panels C and D. **(C)** “Top-view” (membrane distal side) of the cryo-EM map of State2 highlighting the distinct Gβγ binding sites in p110γ and p101 (Gβγ^HD^ and Gβγ^CTD^, respectively). Gβ and Gγ subunits are cyan and green, respectively. **(D)** The corresponding atomic model can be oriented with the geranylgeranyl sites of the two Gβγ subunits and the basic loops of the p110γ C2 domain engaging a common membrane surface.

## Gβγ subunits enage PI3Kγ at two distinct binding sites

To capture snapshots of Gβγ activating PI3Kγ, we purified functional *Sus scrofa* PI3Kγ (p110γ–p101) **(Extended Data Fig 1)** and incubated the protein with a soluble form of human Gβ_1_γ_2_ (Gγ^C68S^) and substrate analogs (ADP·BeF_3_ and C_8_-PIP2, although only ADP was ultimately observed). These samples yielded four independent cryo-EM reconstructions at 3.5-3.9 Å resolution **(Extended Data Figs 2&3; Extended Data Table 1**): two distinct conformations (State1 and 2) of PI3Kγ·ADP in complex with one Gβγ heterodimer bound to the p110γ helical domain (Gβγ^HD^) and another bound to the p101 CTD (Gβγ^CTD^) **(Extended Data Figs 4&5 and Fig 1B-D**), PI3Kγ·ADP in complex with Gβγ^HD^ **(Fig 2A-E)**, and PI3Kγ·ADP alone **(Fig 2F)**. We did not observe any species corresponding to PI3Kγ bound to Gβγ^CTD^ alone, implying that Gβγ^HD^ is the higher affinity site in the absence of a membrane. The Gβγ^HD^ and Gβγ^CTD^ interfaces bury 1600 and 1400 Å^2^, respectively. Gβγ^HD^ and Gβγ^CTD^ bind such that their prenylated Gγ C-termini project towards a plane compatible with the membrane binding loops of the C2 and K domains. Thus State1 and 2 could both represent fully activated PI3Kγ engaging a lipid bilayer **(Fig 1D)**.

**Figure 2.**
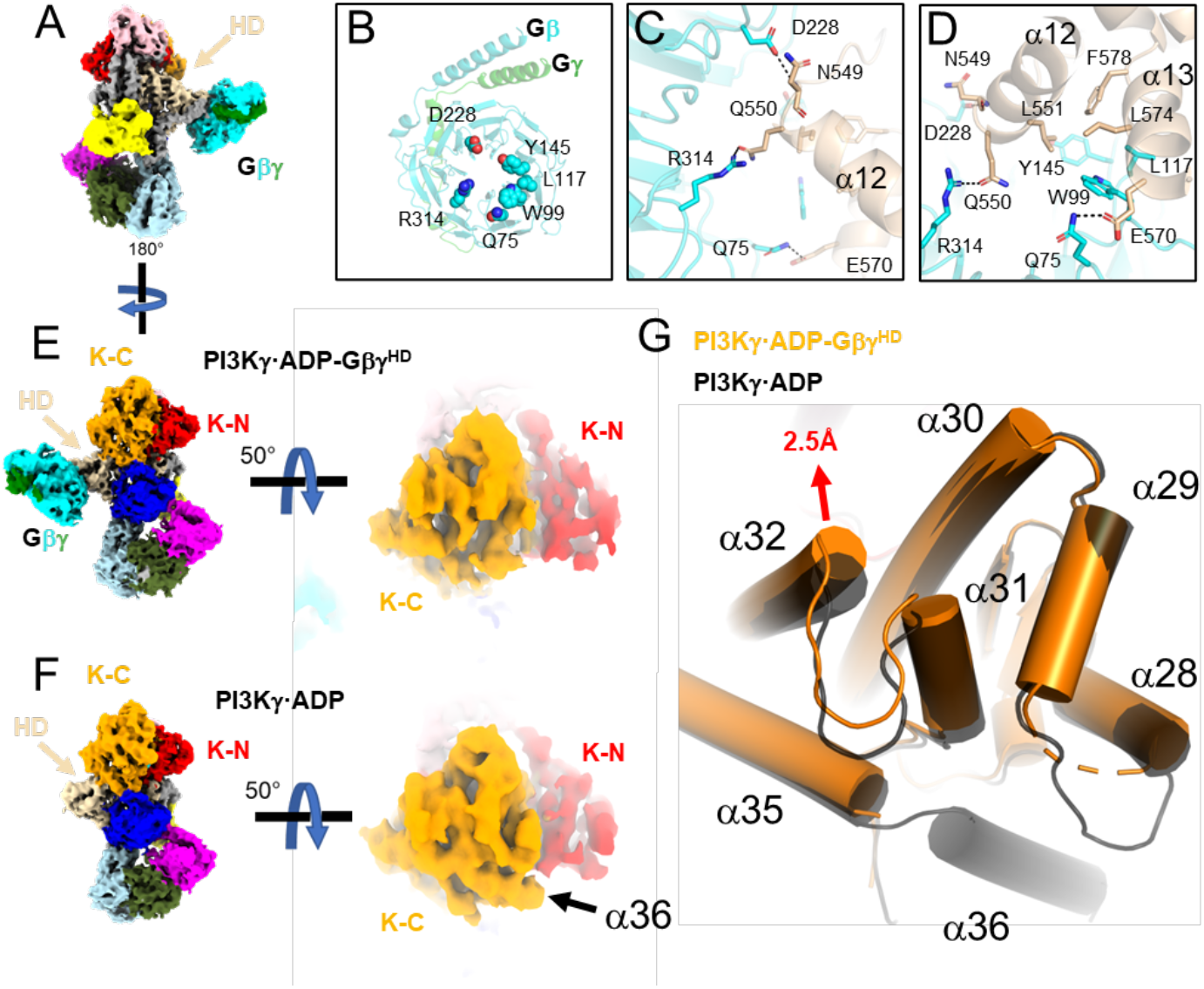
Release of the tryptophan lock and α36 upon Gβγ binding to the p110γ HD. **(A)** Cryo-EM map of PI3Kγ·ADP–Gβγ^HD^ with domains colored as in Fig. 1A. **(B)** Gβγ residues involved in the p110γ helical domain (HD) interaction. **(C-D)** Residues in the Gβγ–HD interface, shown as sticks and colored according to their respective domains. **(E-F)** Comparison of the maps for PI3Kγ·ADP–Gβγ^HD^ and PI3Kγ·ADP. Density for α36 is not observed in PI3Kγ·ADP–Gβγ^HD^. **(G)** Overlay of K-C in PI3Kγ·ADP–Gβγ^HD^ (orange) and PI3Kγ·ADP (black). The movement of the α32 N-terminus is 2.5 Å at the Cα of Leu1004.

In the Gβγ^HD^ interface **(Fig 2B-D, Extended Data Fig 4D-F&5D-F**), residues commonly involved in other Gβγ effector interactions, including Gβ-Ser97, Trp99, and Leu117, engage residues on p110γ α13 (residues 568-578), in particular the side chains of Glu573, Leu574, and His577. In addition, Gβ-Met101, Asp186, Met188, Tyr145, Asp228, Asp246, Arg314, and Trp322 interact with residues at the N-terminus of α12 (residues 548-560), notably Asn549, Gln550, and Leu551 (**Extended Data Fig 4D-F&5D-F**). The side chains of p110γ-Arg552 and Lys553, which were previously predicted to contribute to the Gβγ^HD^ interface^11^, are indirectly involved. The catalytic subunit with p110γ-^552^DD^553^ double mutation, which had significantly reduced sensitivity to Gβγ activation, might have a disrupted interface due to introduction of two helix-destabilizing residues into α12 ^14^. Based on sequence conservation^16^ and the crystal structure of PI3Kβ (PDB entry 2Y3A^17^), we predict that Gβγ could bind the analogous site in the PI3Kβ HD^17^, but not in PI3Kα^18^ or PI3Kδ^19^ (**Extended Data Fig 6**).

Due to the lower quality of density corresponding to the p101 CTD (**Extended Data Figs 4G&5G)**, Alphafold2^20^ was used to guide our modeling of the Gβγ^CTD^ interface. Gβγ^CTD^ binds to a flexible loop of the p101 CTD, which is disordered in structures without bound Gβγ^CTD^. The loop contains a short helix, modeled as residues 709-713, which binds to the “hotspot” of Gβγ^21^. A Gβγ sequence previously implicated in Gβγ binding (^711^KRL^713^) is contained within this element^22^. In our model, the PI3Kγ-Lys711 sidechain electrostatically interacts with Gβ-Asp246, and PI3Kγ-Arg712 protrudes its sidechain into the center of the Gβ WD40 domain to form hydrogen bonds with backbone carbonyls of Ser155 and Met188, and PI3Kγ-Leu713 forms hydrophobic interactions with Gβ-Trp99 and Met101 **(Extended Data Figs 4J-L&5J-L)**. An extended ribbon formed by the β8 and β9 strands of the CTD may form a secondary contact with a hydrophobic patch on Gβγ^CTD^ near residues Leu55 and Phe335, but it was not possible to model the density. This unusual loop contains conserved basic and a few conspicuously exposed hydrophobic sidechains (*e.g.* Phe789) that may form stabilizing interactions with the plasma membrane upon binding of Gβγ^CTD^. The basic electrostatic surface created by this loop is continuous with that of the membrane-proximal side of Gβγ^CTD^. This may help explain why in cells that the p101 site is more strongly recruited to the plasma membrane than p110γ ^13^.

States 1 and 2 of the PI3Kγ·ADP–Gβγ^HD^–Gβγ^CTD^ complex primarily differ in the orientation of their p101 CTDs relative to the rest of the protein. Their CTD domains are related by a ∼ 7° rotation around the CD and a translation of ∼3 Å. Consequently, the two Gβγ^CTD^ domain centers translate ∼15 Å. Notably, a β4-β5 hairpin loop that extends from the central sheet of the p101 CTD domain and contacts the C2 domain (**Fig. 1D, 3C**), although involving a relatively small amount of surface area, is maintained in all the various structures/conformations we determined. This coupling may transmit membrane/Gβγ binding events and conformational changes occurring with the CTD with the membrane binding loops of the adjacent C2 domain and could be a route of overall allosteric control.

## Conformational changes induced by Gβγ binding

For comparison with the Gβγ complex data sets, we generated a 3 Å reconstruction of ligand-free PI3Kγ (**Extended Data Table 1, Fig 3, Extended Data Fig 7**). We also determined reconstructions of the PI3Kγ· ADP and PI3Kγ·ATP complexes at resolutions of 3.9 and 3.3 Å, respectively (**Extended Data Table 1, Extended Data Figs 2-3 and 8-9**). These structures are nearly identical to that of PI3Kγ alone (RMSD of 0.54 Å for 1378 Cα atoms comparing PI3Kγ· ADP to PI3Kγ; RMSD of 0.62 Å for 1285 Cα atoms comparing PI3Kγ·ATP to PI3Kγ; RMSD of 0.65 Å for 1330 Cα atoms comparing PI3Kγ·ADP to PI3Kγ·ATP), suggesting negligible conformational changes occur in PI3Kγ upon binding nucleotides in the active site. Density for the K-C C-terminal helix α36 is observed in all these cryo-EM structures **(Fig 2F, Fig 3D, Extended Data Fig 9B)**, consistent with ATP or ADP binding not disturbing the structure in the kinase C-lobe and or releasing the tryptophan lock ^23^ **(Extended Data Fig 10)**.

**Figure 3.**
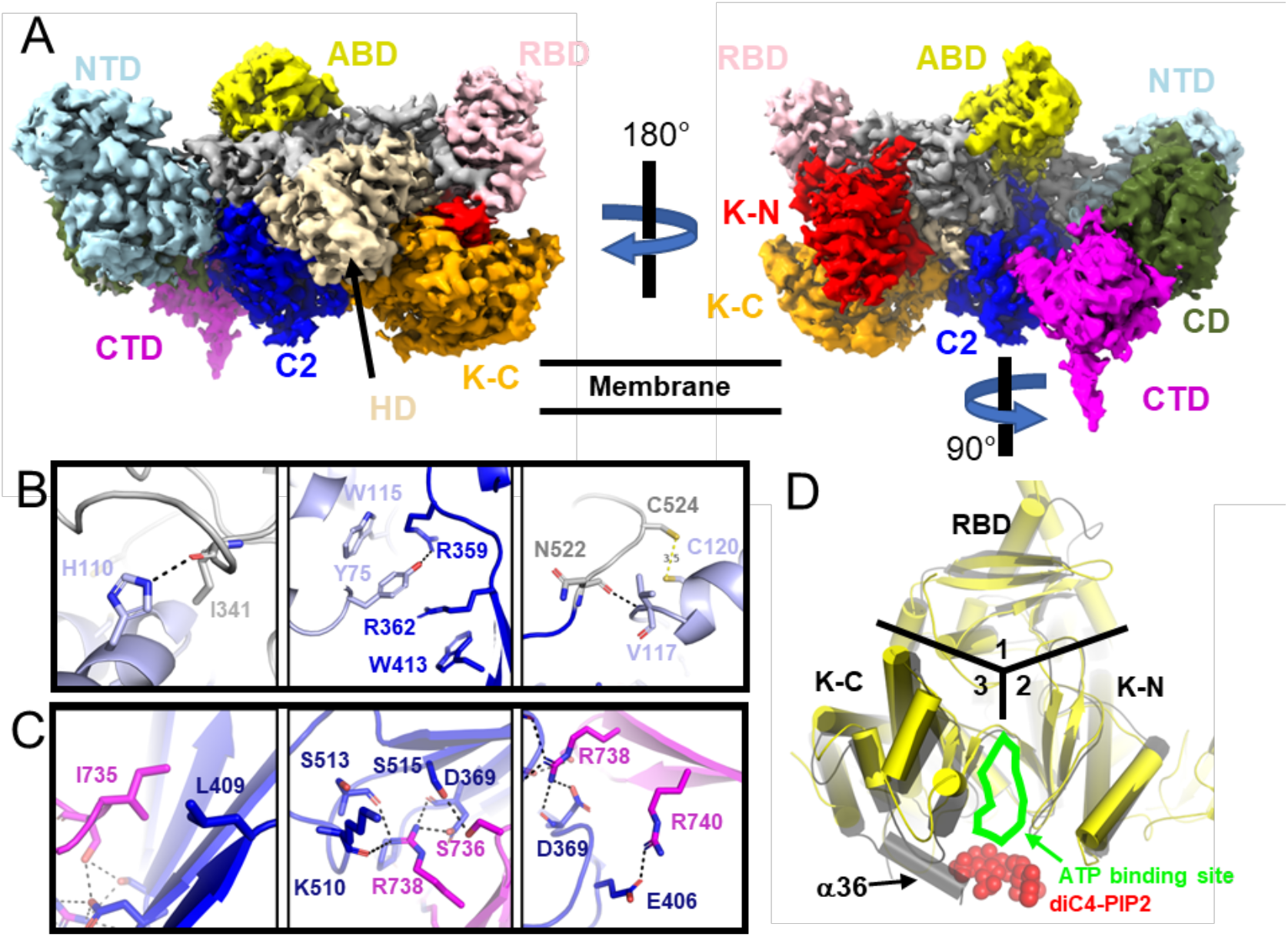
Cryo-EM structure of PI3Kγ. **(A)** Composite map (also see the **Extended Data Fig 2H**) colored as described in **Fig 1A**. The NTD **(B)** and a hairpin loop in CTD **(C)** of p101 form distinct contact sites with the C2 domain and its preceding linker in p110γ. The side chains of interacting residues in the p110γ RBD-C2 linker (grey), p110γ C2 (blue), p101 NTD (light blue), and p101 CTD (magenta) are shown as sticks. Hydrogen bonds are shown as dashed lines. **(D)** Superimposition of PI3Kγ with the crystal structure of PI3Kα·diC4-PIP2. RBD, K-N, and K-C domains are located in areas 1, 2, and 3, respectively. The ATP binding site is indicated, and red spheres show the diC4-PIP2.

In the PI3Kγ·ADP–Gβγ^HD^ structure, the linker between the C2 and HD (residues 522-545) is better ordered than in the PI3Kγ structure. However, the C-terminal helix α36 (1081-1092) in K-C becomes disordered relative to the PI3Kγ·ADP structure **(Fig 2E-F)**. Displacement of α36 has been proposed to require the release of the tryptophan lock, in which Trp1080 loses its contacts with Pro989 on α31, Phe902 on the α28-α29 loop (residues 894-905), and Asp904 (**Extended Data Fig 10A**). In PI3Kγ·ADP–Gβγ^HD^, α31 and α32 shift, and the α28-α29 loop becomes disordered (**Fig 2G, Fig 4A-B, Extended Data Fig 10D**). Therefore, binding of Gβγ^HD^ results in conformational changes in K-C, even in the absence of a lipid membrane or lipid substrates. The packing of a conserved residue, Leu564, in the α12-α13 loop of the HD is perturbed upon Gβγ^HD^ binding (**Fig 4C**) such that it loses buried surface area with helices α33 and α34 of the K-C **(Fig 4C)**, which connect to the “α31, α32, α35” bundle (**Extended Data Figs 10E-H**). Reorganization of this contact may be the allosteric conduit by which Gβγ^HD^ binding opens the kinase active site. Intriguingly, we observed higher movement of helices α28, α29, α30, α31, α32, α35 in K-C upon binding of Gβγ^CTD^ in the State1 and 2 reconstructions (**Fig 4B**). This suggests that Gβγ^HD^ binding may only lead to partial kinase activation (**Fig 4B**).

**Figure 4.**
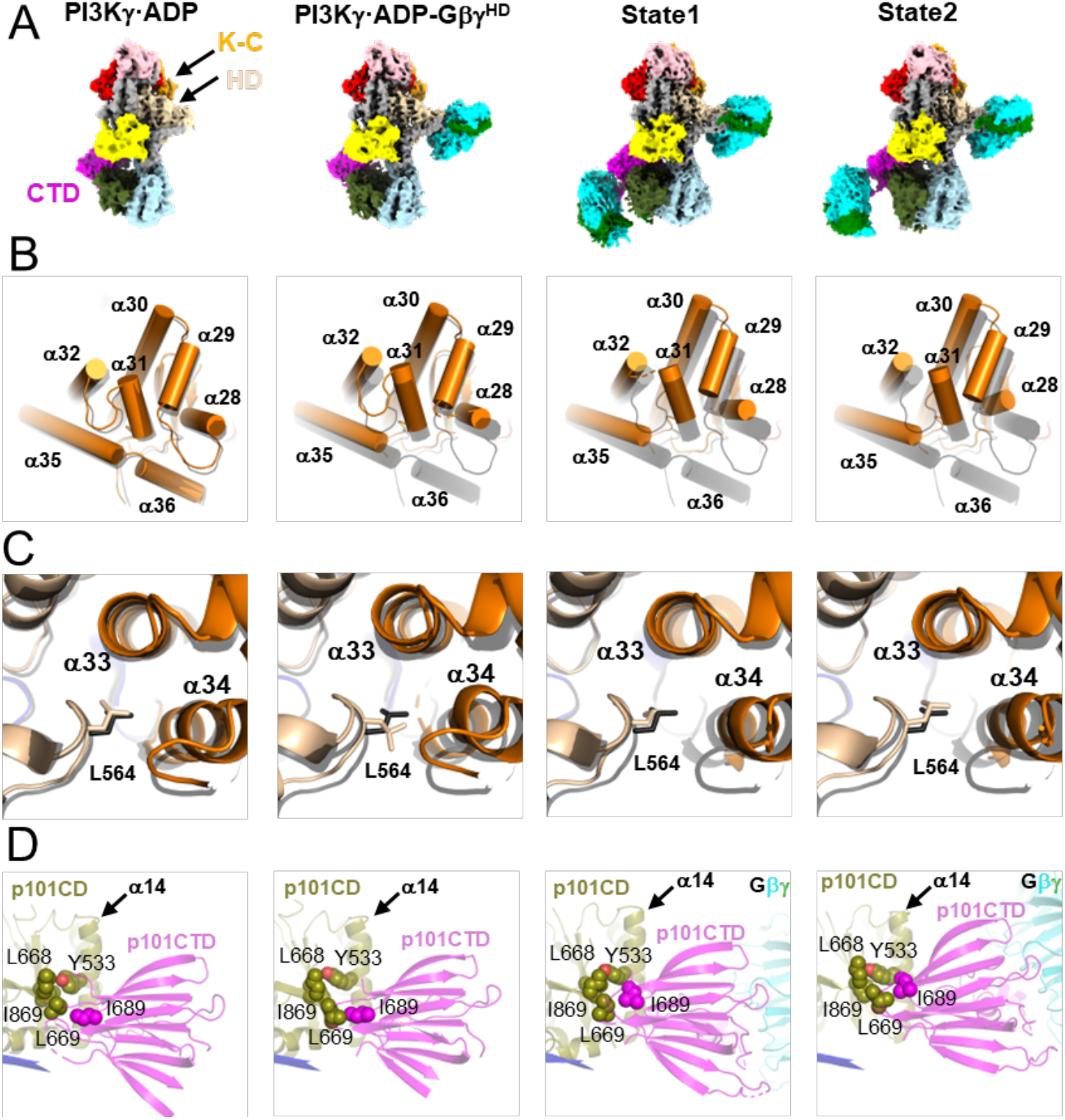
Structural changes in PI3Kγ as a function of Gβγ binding. **(A)** Cryo-EM maps of PI3Kγ·ADP, PI3Kγ·ADP–Gβγ^HD^, and States 1 and 2 of PI3Kγ·ADP–Gβγ^HD^–Gβγ^CTD^. **(B)** Comparison of helices in K-C. **(C)** Comparison of Leu564 sidechain packing. In **(B-C)**, the structures of PI3Kγ·ADP, PI3Kγ·ADP–Gβγ^HD^, State 1 and 2 of PI3Kγ·ADP–Gβγ^HD^–Gβγ^CTD^ were individually superimposed with that of PI3Kγ (black). **(D)** Variability in the position of the p101 CTD. Interacting residues, Tyr533, Leu668, Leu669, and Ile869 in the p101 CD and Ile689 in the p101 CTD are shown as spheres.

Upon Gβγ^CTD^ binding, the p101 CTD changes position relative to all the other domains of PI3Kγ, and it is also different between states 1 and 2 of PI3Kγ·ADP–Gβγ^HD^–Gβγ^CTD^. A conspicuous marker of this conformational variability is how the side chain of Ile689 residue packs against the α14 helix of CD (**Fig 4D**), suggesting that this residue is important for allosteric coupling. Residues 816-830 of p101-CTD form more extensive interactions with the CD in the Gβγ^CTD^-bound complexes relative to PI3Kγ alone, consistent with previously observed HDX protection of this region upon Gβγ binding (Vadas, 2013).

### Functional consequences of Gβγ binding

Based on the structural results we hypothesized that the binding of Gβγ^HD^ or Gβγ^CTD^ allosterically triggers conformational changes in the K-C that facilitate catalysis **(Fig 4, Fig 5A)**. To validate each observed binding site, we mutated key residues in each site: p110γ-L551A, p101-^711^Lys-Arg^712^ to ^711^Gly-Ser^712^ (CTD-GS), and p101-^711^Lys-Arg-Leu^713^ to ^711^Gly-Ser-Ser^713^ (CTD-GSS) **(Fig 5B)**. We also generated L564S and I689T variants **(Fig 5B)** to probe the importance of allosteric transitions we observed to be mediated by the binding of Gβγ^HD^ and Gβγ^CTD^, respectively. We then compared the specific enzyme activity of each variant to that of wild-type (WT) PI3Kγ at varying concentrations of Gβγ using liposome-based kinase assays (**Fig. 5C-F, Extended Data Table 3**). Disruption of either Gβγ binding site significantly impaired enzyme activity **(Fig. 5C-D)**. The L564S mutation resulted in a total loss of Gβγ-mediated activation **(Fig. 5E)**, consistent with it playing critical role in propagating signals from both Gβγ binding sites. This position is conserved as Leu547 in PI3Kβ **(Extended Data Fig 6B)**. Significantly decreased activity in the dose-response curve for the p101^I689T^ mutant **(Fig 5F)** likely reflects less efficient coupling between the p101-CTD and CD (**Figs 1D&4D**).

**Figure 5.**
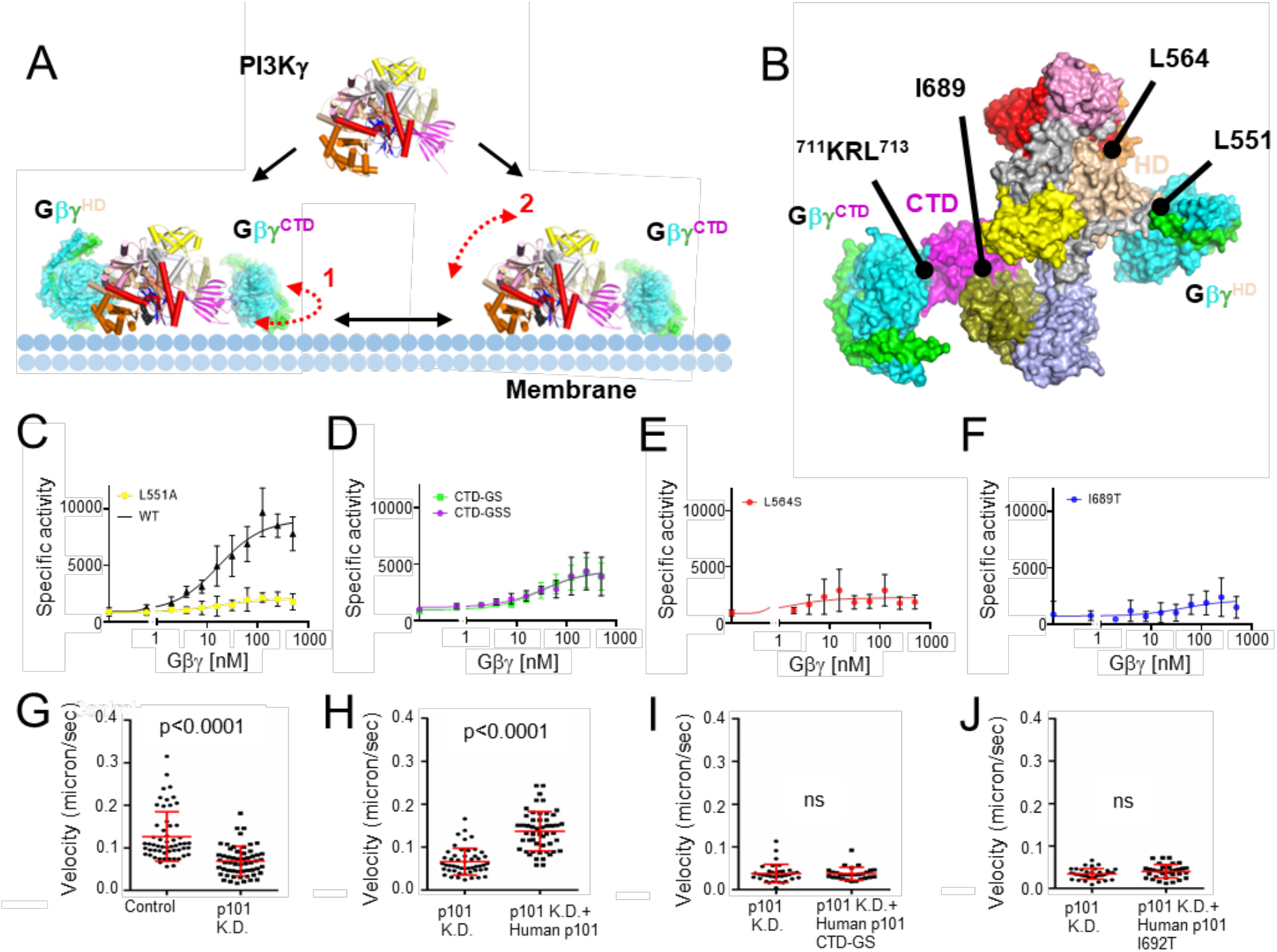
Functional characterization of Gβγ-mediated PI3Kγ activation. **(A)** PI3Kγ membrane recruitment via two Gβγ subunits, Gβγ^HD^ and Gβγ^CTD^. Dashed red line 1 indicates the conformational flexibility of the CTD domain of PI3Kγ upon Gβγ^CTD^ binding that would allow it to move along the plane of the membrane, as suggested by the State 1 and 2 structures we determined. Dashed red line 2 represents the fact that the p110γ subunit is still not fully engaged with the membrane even when tethered to the membrane (in this example by Gβγ^CTD^), as suggested previously^13^. **(B)** Surface representation of the PI3Kγ·ADP–Gβγ^HD^–Gβγ^CTD^ State 2 structure. The sites targeted by site-directed mutagenesis represent those for Gβγ binding (p110γ-Leu551 and p101-^711^KRL^713^) and regulation (p110γ-Leu564 and p101-Ile689). **(C-F)** Dose response curves of PI3Kγ variants (WT, black triangles; p110γ−L551Ap101, yellow circles; p110γp101-CTD-GS and p110γp101-CTD-GSS, green and purple circles; p110γ−L64Sp101, red circles; p110γp101-I689T, blue circles. Specific activity is nmol ADP·μmol enzyme^-1^·min^-1^. **(G-J)** Neutrophil migration imaging and tracking in a zebrafish embryo 3 days post-fertilization. **(G)** Quantification of neutrophil tracks under control (PBS) or p101 knock down (MO) background. Representative images are shown in **Extended Data Fig 11A-B. (H-J)** Quantification of neutrophil tracks under p101 knockdown (MO) background of neutrophils without or with expressing human WT p101 **(H)**, mutant human p101-CTD-GS **(I)**, or mutant human p101-I692T **(J)**. Representative images are shown in **Extended Data Fig 11C-E**. In **(G-J)**, three independent experiments with n>20 are compiled. p values represent overall effects using the Gehan-Breslow-Wilcoxon test.

To further examine regulatory functions of the Gβγ^CTD^ site of p101 on PI3Kγ activation, we created morpholino p101 knock-down zebrafish animals **(Fig 5G-J, Extended Data Fig 11, and Movies 1, 2)** and measured neutrophil motility either without **(Fig 5G)** or with the expression of WT **(Fig 5H)** or mutant human p101 proteins: human p101 CTD-GS **(Fig 5I)** and human p101 I692T (equivalent to I689T in *Sus scrofa*) **(Fig 5J)**. WT human p101 **(Fig 5H)** rescued neutrophil motility and their resulting behavior was comparable to normal zebrafish neutrophils **(Fig 5G)**. However, with the mutant human p101 variants neutrophil motility remained impaired (**Figs. 5H-J**). These results suggest that the p101 regulatory subunit not only functions to recruit PI3Kγ to the plasma membrane but also activates PI3Kγ allosterically because the I692T mutant should not be impaired in binding membrane-associated Gβγ **(Fig 4D)**, yet still fails to rescue. We could not rescue p110γ knock-out zebrafish model animals with human WT p110γ, which prevented us from testing mutations at the Gβγ^HD^ site.

## Discussion

HDX-MS and other biochemical studies have previously provided insights into possible Gβγ binding interfaces on PI3Kγ and regions that potentially undergo conformational changes, such as residues 579-607 in the p110γ helical domain, helices α31 (989-995), α35 (1060-1078), and α36 (1085-1090) in the p110γ K-C, and the p101 CTD. Because these same regions also showed perturbed HDX signals when PI3Kγ attaches to the liposome vesicles, a model of membrane-induced conformational change was proposed to explain activation by Gβγ ^11,14^. Based our data, we propose that in addition to any effects the lipid bilayer might have, soluble (unprenylated) Gβγ can itself trigger an allosteric transition involving p110γ-Leu564, which disseminates the Gβγ-mediated activation signal through helices α32 and α31 to the α28-α29 loop, leading to an opening of the tryptophan lock and release of α36 blocking the active site of the kinase domain (**Figs 2G, 4B, Extended Data Fig 10**). Comparing our cryo-EM structure of PI3Kγ and PI3Kγ-ATP with the crystal structure of PI3Kα bound to diC4-PIP2 (PDB entry 4OVV), the PIP2 headgroup would clash with α36 in both PI3Kγ (**Fig 3D**) and PI3Kγ-ATP (**Extended Data Fig 9B**). Consistent with this idea, α36 was not ordered in the PI3Kα crystal structure. The binding of Gβγ^CTD^ alters the interface between the p101-CTD and CD domains with the assistance of Ile689 and also likely pulls on the C2 domain via the β4-β5 hairpin loop (**Fig 1D&4D**). Although the long-range allosteric pathway to the kinase domain is less clear, Gβγ^CTD^ binding leads to an even greater displacement of helices in the K-C in the PI3Kγ·ADP–Gβγ^HD^–Gβγ^CTD^ complexes (**Fig 4D**). We have yet been unable to isolate PI3Kγ in complex with Gβγ^CTD^ alone, which suggests that either it is dependent on the binding of Gβγ^HD^ in our system, or, trivially, that it is too disordered for the processing programs to reconstruct in the absence of Gβγ^HD^. Because both the p110γ-L564S and p101-I689T mutations individually nearly eradicate activation by Gβγ, it implies these regions are involved in a shared allosteric network necessary for full activation. The residual activity of these variants in our assays could then reflect activation via membrane recruitment. Finally, because perturbation of the Gβγ^CTD^ site does not reduce activation to the level of p110γ-L551A, it implies that the HD is the dominant site involved in allosteric regulation, a conclusion that has been suggested previously in cell based data^13^.In the crystal structure of p110γ–H-Ras·GDPPNP (PDB entry 1HE8)^24^, similar helical movements in K-C were observed **(Extended Data Fig 12A-D)**, suggesting that both Ras and Gβγ binding can release the tryptophan lock and promote PIP2 binding by allosterically tuning the surrounding helices (**Fig 4B**).

In summary, we have described cryo-EM structures of PI3Kγ in various ligand/Gβγ protein-bound states that provide new insights into mechanisms underlying Gβγ-mediated activation of PI3Kγ (**Fig 4 & Fig 6**), and which parallel those of the Ras binding to the RBD (**Fig 6 and Extended Data 12A-D)**. There likely exist other paths for PI3Kγ activation as evidenced by cancer-related somatic mutations, such as E347K in the RBD-C2 linker (residues 311-357) found in pancreatic ductal carcinoma and large intestine adenocarcinoma^25^, and R472C in the C2 domain found in prostate cancer^25^, which show enhanced Gβγ-mediated activation in *in vitro* assays^11^.

**Figure 6.**
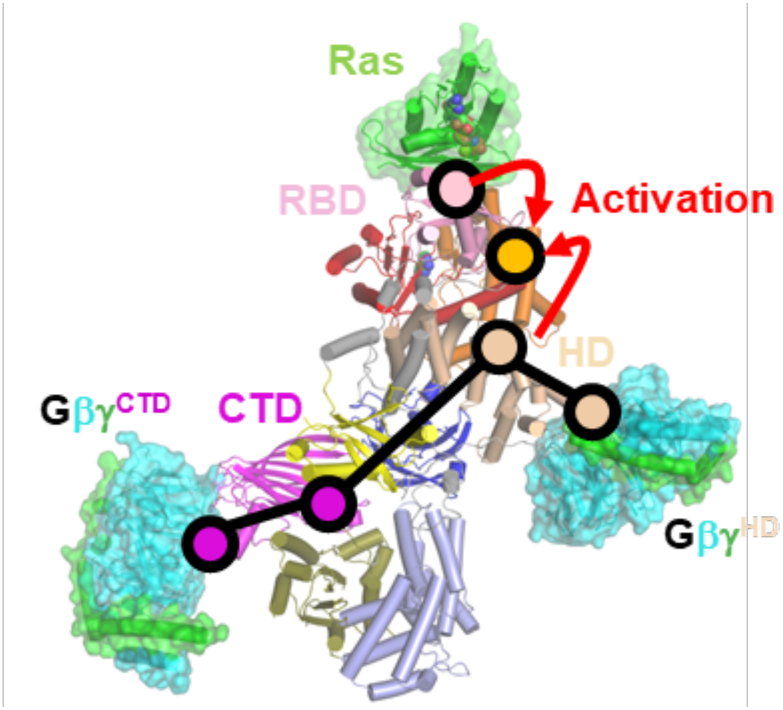
Proposed allosteric pathways in PI3Kγ. Purple circles indicate domains proposed to be involved in allosteric activation by Gβγ^CTD^. Wheat circles indicate key regions involved in allosteric activation by Gβγ^HD^. Based on our data, the two networks would seem to converge on the interface between the HD and K-C domain. The psink circle indicates the Ras binding interface with RBD, demarking the route for allosteric activation by Ras. The orange circle demarks the kinase active site. Note that the NTD and CD of p101 seem to form a rigid scaffold with the C2, HD, and K-N domains of p110γ, providing a framework within which the p101 CTD and K-C domains exhibit dynamics essential for allosteric control of PIP3 production.

## Methods

### Expression constructs and purification of PI3Kγ variants

Full-length p110γ and p101 from *Sus scrofa* were inserted into the pFastBac™ Dual vector (Thermo Fisher) for use in the Bac-to-Bac Baculovirus Expression System (used without authentication or testing for mycoplasma). A His_6_-tag was added to the N terminus of p110γ, and a FLAG-tag to the C terminus of p101. Mutations in p110γ and p101 were created using the overlap PCR technique **(Extended Data Table 2)**, followed by restriction enzyme digestion and ligation (*Eco*RI and XbaI for p110γ, and *Nhe*I and *Kpn*I for p101). Sf9 insect cells were infected by baculovirus for each PI3Kγ variant for two days and harvested (6,000xg for 15 minutes). For purification, freshly prepared or thawed cell pellets were resuspended in cold lysis buffer containing buffer A (20 mM HEPES pH 8.0, 100 mM NaCl) and 20 mM EDTA, 0.1 mM each of PMSF, leupeptin (10 μM) and lima bean trypsin protease inhibitor (0.1 mg/ml), and the SIGMAFAST™ protease inhibitor cocktail (1 tablet in 100 ml buffer). The cells were resuspended and then lysed using an Avestin C3 emulsifier. The lysed cells were centrifuged in a Beckman JA-25.5 rotor at 30,000xg for 20 min. A 40 ml supernatant was then rocked with ∼80 μl anti-FLAG magnetic beads (pre-equilibrated with lysis buffer) for at least 2 hours. The anti-FLAG magnetic beads bound with proteins were washed with 10 ml lysis buffer for 5 times, followed by elution with 50 μl buffer A with Flag peptide at a final concentration of 1 mM. We assessed the purity and the quantity of PI3Kγ proteins via SDS-PAGE. Freshly purified protein samples were used immediately for cryo-EM and biochemical studies.

### Expression and Purification of WT (Gβγ^WT^) and soluble Gβγ (sGβγ^C68S^)

The WT human geranylgeranylated Gβ_1_γ_2_ protein (Gβγ^WT^) with an N-terminal His_6_ tag on Gβ was expressed using the Sf9 expression system and purified by Ni-NTA affinity chromatography as described previously^4^. Briefly, the cells were infected by baculovirus for two days and harvested (6,000xg for 15 minutes). The cell pellet was resuspended with Buffer A (20 mM HEPES pH 8, 100 mM NaCl, and 1 mM MgCl_2_) and lysed by a high-pressure homogenizer (Emulsiflex C3). The cell debris was pelleted using centrifugation (30,000xg for 15 minutes). The membrane fraction in the supernatant was pelleted by ultracentrifugation (186,000xg for 40 mins), followed by solubilization with Buffer A supplemented with 1 % sodium cholate. After another run of ultracentrifugation, the supernatant containing Gβγ^WT^ was loaded onto Ni^2+^ resin (HisPurTM Ni-NTA resin, ThermoFisher) equilibrated with Buffer A. Impurities were removed from the resin during the wash step with 20 column volumes of Buffer B (20 mM HEPES pH 8, 100 mM NaCl, 1 mM MgCl_2_, 10 mM CHAPS, and 10 mM imidazole pH 8), and the Gβγ protein was eluted with 5 column volumes of Buffer C (20 mM HEPES pH 8, 100 mM NaCl, 1 mM MgCl_2_, 10 mM CHAPS, and 150 mM imidazole pH 8). The eluted protein was dialyzed against Buffer D (20 mM HEPES pH 8, 0.5 mM EDTA, 2 mM MgCl_2_, 10 mM CHAPS, and 1 mM DTT) and loaded onto an anion exchange column (Q column). The column was first washed with Buffer D until the UV280 signal was stable. Gβγ^WT^ was eluted with a linear gradient to 30 % Buffer E (20 mM HEPES pH 8, 1 M NaCl, 0.5 mM EDTA, 2 mM MgCl_2_, 10 mM CHAPS, and 1 mM DTT). The eluted fractions were analyzed by SDS-PAGE and further purified by size exclusion chromatography (SEC) using Buffer F (20 mM HEPES pH 8, 100 mM NaCl, 0.5 mM EDTA, 2 mM MgCl_2_, 10 mM CHAPS, and 1 mM DTT). Peak fractions from SEC were pooled, fresh-frozen in liquid nitrogen, and stored at -80°C. The expression and purification steps for the soluble form of Gβγ (sGβγ^C68S^) are similar to that for Gβγ^WT^, albeit the steps for collecting membrane fraction by ultracentrifugation were excluded, and no detergents (cholate, CHAPS) were added to the buffer solutions^26^.

### PI3K Assay

The ADP-Glo lipid kinase assay (Promega) was used to monitor ADP production, which corresponds to PIP3 production, following the manufacturer’s protocol. Liposomes containing phosphatidylserine and PIP2 (3:1 ratio mol/mol) were purchased (Promega) and prepared to a final concentration of PIP2 at 25 μM. A varied concentrations of Gβγ^WT^ (500, 250, 125, 62.5, 31.25, 15.6, 7.8, 3.9, 1.95, 0.978 and 0 nM) were added to the liposomes, followed by addition of the WT or mutant PI3Kγ proteins (1-10 nM) with ATP (final concentration of 100 μM). Reactions were performed at room temperature for 40 minutes, followed by the addition of the ADP-Glo reagent for 40 minutes and the addition of the fluorescent substrate. The specific activity of each variant is summarized in **Extended Data Table 3** [N=8∼14, 3 parameter curve fitting using GraphPad Prism 9].

### Cryo-EM sample preparation and data collection and processing

Cryo-EM samples consisted of PI3Kγ (0.05-0.2 mg/ml), PI3Kγ supplemented with 2 mM of either ATPor ADP·BeF_3_^27^, 2 mM C_8_-PIP2, and sGβγ^C68S^ (0.2 mg/ml)^28^ were prepared for data collection. The UltrAuFoil gold grids (R1.2/1.3, 300 mesh) were glow-discharged (25 mA, 60 seconds) and soaked into 100 mM MgCl_2_, followed by applying 3.5 μl of protein samples onto the grids before plunge-freezing using the Vitrobot Mark IV system (Thermo Fisher Scientific)(blotting force=2, blotting time=3.5 seconds, wait time= 5 minutes). All datasets were collected using a Titan Krios cryo-electron microscope, and micrographs were recorded as movies on the Gatan K3 detector (nominal magnification: 81,000x, magnified pixel size: 0.54 Å in super-resolution mode). The total electron dose for data collection was ∼ 56 e^-^/A^2^. Motion correction to align the movie frames for each of the data sets was performed using motioncor2 software^29^ implemented within CryoSPARC. The CTF of each aligned micrograph was estimated using the CTFFind4 module of CryoSPARC. Particle picking, 2D classification, initial model generation, 3D classification, and 3D refinement were also performed in CryoSPARC. Each dataset was split into two independent sub-datasets and then subjected to individual data processing for map reconstructions and resolution estimation. The nominal resolution was determined based on a Fourier shell correlation cutoff of 0.143^30^. Statistics for all datasets are summarized in **Extended Data Table 1**. For processing the data of PI3Kγ supplemented with ADP·BeF_3_ and sGβγ^C68S^, we used RELION 3.1^31^ for additional 3D classification during the process **(Extended Data Fig 2)**. In the case of PI3Kγ (**Extended Data Fig 7**), there was a well-resolved p110γ density but relatively low-resolution (3.5∼4.5Å) p101 density (**Extended Data Fig 7E**). The density in the N-terminal and C-terminal halves of p101 was improved by local refinement with different fulcrum points and varying sizes of mask applied (**Extended Data Fig 7F-G**). The locally refined maps were combined to generate a composite map (**Fig 3A, Extended Data Fig 7H-I)** (phenix_combine.focus.maps)^32^ for model building.

### Structure modeling and refinement

Model building was performed using Coot^33^, and alternated with real-space structure refinement using phenix.real_space_refine in Phenix^32^ with refinement strategies of global minimization (minimization_global) and B-factor refinement (adp). Because the p101 structure was unknown at the time the first maps were generated, the NTD and CD of the protein were built *ab initio* into locally refined maps. The advent of AlphaFold2 contributed to predicting the structure of p101, helped verify our modeling, and allowed us to build the p101 CTD density. During this time, the structure of PI3Kγ (human p110γ, pig p101) in complex with a nanobody (PI3Kγ-Nb) that binds the p101 CTD became available and was also assessed for consistency (RMSD of 1.34 Å for 972 Cα atoms in p110γ; and RMSD of 2.9 Å 501 Cα atoms in p101). As indicated by the high RMSDs, the structures have significant differences, most notably in the registration of several secondary structure elements in the p101 NTD and CD, and in regions of the ABD of p110γ, as well as in the topological modeling of the p101 CTD (**Extended Data Fig 13A-B)**. The bound nanobody also seems to alter the orientation of the CTD^11^. Thus, we used our base atomic model of PI3Kγ as the seed for all the other reconstructions. The initial models of the G ^HD^ and G ^CTD^ complexes were generated by AlphaFold multimer^20^, and the resulting G -bound p110 helical domain and the p101 CTD were then merged with the rest of the PI3K model to set up MDFF procedures^34-36^. The grid scaling (gscale) value was set to 0.5. Suggested restraints regarding secondary structures, peptide bonds, and chirality also were applied during MDFF fitting ^35,36^. For State1 and 2 of PI3K ·ADP–G ^HD^–G ^CTD^, the MDFF simulation was performed for 300 picoseconds until convergence of the model RMSD^36^ (**Extended Data Fig 14A&C**), followed by 3000 steps (3 picoseconds) of energy minimization. The quality of the MDFF fittings was estimated using the CCC generated by VMD ^36^ (**Extended Data Fig 14B&D**). All models were subjected to rounds of cryo-EM real-space refinement in Phenix^32^ alternating with manual adjustments using Coot. DAQ-score^37^ was used to validate final models against the 3D reconstruction data, with an example shown in **Extended Data Fig 15**. Results are summarized in **Extended Data Table 1**.

### Morpholino oligonucleotides (MO) microinjection

Splice morpholino for pik3r5-202 e2i2 (5’-TACAAACATTCTCACCTCTCTGGCT-3’) was purchased from GeneTools, LLC, and resuspended in distilled water. Stocks are stored at RT with a concentration of 1 mM. For the knock-down experiments, one nanoliter of MO was injected into the yolk of 1-cell stage zebrafish embryos at 0.2 mM concentration. IThe human PIK3R5 sequence was cloned from Plasmid #70463, R777-E179 Hs.PIK3R5 (Addgene) into the *tol2* transposon system with *lyzC* promoter for neutrophil-specific expression^38^. GFP with 2A peptide was inserted to generate the Tol2-lyzC-GFP-2A-PIK3R5 (p101) plasmid. For the following rescue experiments, microinjections of zebrafish embryos were performed by injecting 1 nL of a mixture containing 25 ng/μL plasmid and 35 ng/μL Tol2 transposase mRNA in an isotonic solution into the cytoplasm of embryos at the 1-cell stage. Mutations were generated using primers that bind to human PIK3R5 sequences: 5’-GATCTTCACCCACTCCTTGGAGCTGGGTCAC-3’ and 5’-GAGTGGGTGAAGATCTCTGTCGTCTTCTCCCC-3’ for the I692T mutation; 5’-GCAGCGGGTCGCTGGGCATCGATGGCGACC-3’ and 5’-CCAGCGACCCGCTGCCAGGACCTGACGC-3’ for the p101 CTD-GS mutation.

### Live imaging

Larvae at 3 days post fertilization (dpf) were mounted with 1% low melting agarose and 0.02% tricaine on a glass-bottom dish, and imaging was performed at 28°C. Time-lapse fluorescence images in the head mesenchyme were acquired with 10x magnification on BioTek Lionheart FX Automated Microscope (Agilent). The green and red channels were acquired sequentially with 488-nm laser 561-nm laser, respectively. Videos of neutrophil motility were taken with stack slices less than 10 μm. The fluorescent stacks were flattened using the maximum intensity projection and overlayed with each other for representation. MtrackJ in ImageJ was used for tracking neutrophil motility.

### Data Availability

All data needed to evaluate the conclusions in the paper are presented in the paper and/or the Supplementary Materials. Additional data related to this paper are available upon reasonable request from the authors. The structures of PI3K, PI3K ·ATP, PI3K ·ADP, PI3K ·ADP–G ^HD^, State1 and 2 of PI3K ·ADP–G ^HD^–G ^CTD^, as well as their associated cryo-EM reconstructions, have been deposited into the Protein Data Bank under accession codes 8SO9, 8SOA, 8SOB, 8SOC, 8SOD and 8SOE, and the Electron Microscopy Data Bank under accession codes EMD-40650, EMD-40651, EMD-40652, EMD-40653, EMD-40654, and EMD-40655, respectively.

### Statistics

I*n vitro* and *in vivo* assays were conducted with at least three technical replicates (n=3). The dose response curves were fitted to the *in vitro* lipid kinase assays with a three-parameter logistic nonlinear regression model implemented in GraphPad Prism 9. For the Zebrafish K.D. and rescue experiments, three independent experiments with n>20 are compiled. The p values represent overall effects using the Gehan–Breslow– Wilcoxon test.

## Author Contributions

C.-L.C. and J.J.G.T. conceptualized the study. S.K.R. created the WT PI3K construct and C.-L.C. and Y.-C.Y. created the mutant PI3K constructs. C.-L.C. produced and purified WT and mutant PI3K proteins for conducting lipid kinase assays. Y.-C.Y. produced the WT G proteins. C.-L.C. and T.K. collected cryo-EM data, and C.-L.C. performed cryo-EM data analysis. C.-L.C. performed molecular dynamics flexible fitting. C.-L.C. and J.J.G.T. performed atomic modeling for all the cryo-EM reconstructions of PI3K without or bound with ligand, and G. R.S and Q.D. conducted the morpholino knock-down and rescue experiments using the zebrafish model system. C.-L.C. and J.J.G.T. wrote the original draft, and all authors further edited the manuscript.

## Acknowledgement

The authors acknowledge funding from NIH grants CA254402 (J.J.G.T), CA221289 (J.J.G.T.), CA023168 (J.J.G.T.), HL071818 (J.J.G.T.), R35GM119787 (Q.D.), American Heart Association Postdoctoral fellowship (#834497, C.-L.C.), Purdue Center for Cancer Research (PCCR) Pilot Grant 2020-21 Cycle 2, PCCR Shared Resource 2020-2021 Cycle 1 (Grant #30001612), the Walther Cancer Foundation (J.J.G.T.) and the technical support of Purdue Life Sciences Electron Microscopy Facility for cryo-EM data collection.

## Notes

### Competing Interest Statement

The authors have declared no competing interest.

