## Supplementary Data for "Molecular basis for Gβγ-mediated activation of phosphoinositide 3-kinase γ"


**Extended Data includes:**

Extended Data Figs 1-15

Extended Data Tables 1-3

Extended Data Movies 1-2

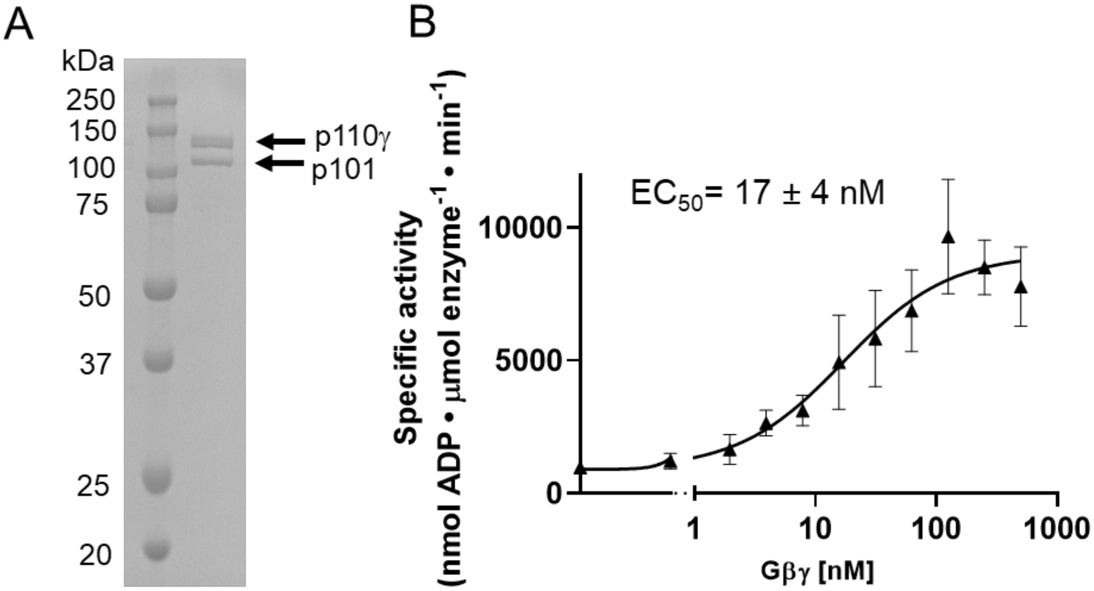


**Extended Data Figure 1. Purification of functional PI3Kγ (A)** SDS-PAGE analysis of PI3Kγ. The protein was purified via anti-FLAG affinity pull-down and eluted from the beads with 3X-FLAG peptides at a final concentration of 20 μM. **(B)** Activation of PI3Kγ by Gβγ. PI3Kγ activity measured at varying concentrations of Gβγ ([Gβγ]) from 0 to 500 nM. [n=8; error bars represent SD] (see Extended Data Table 3).

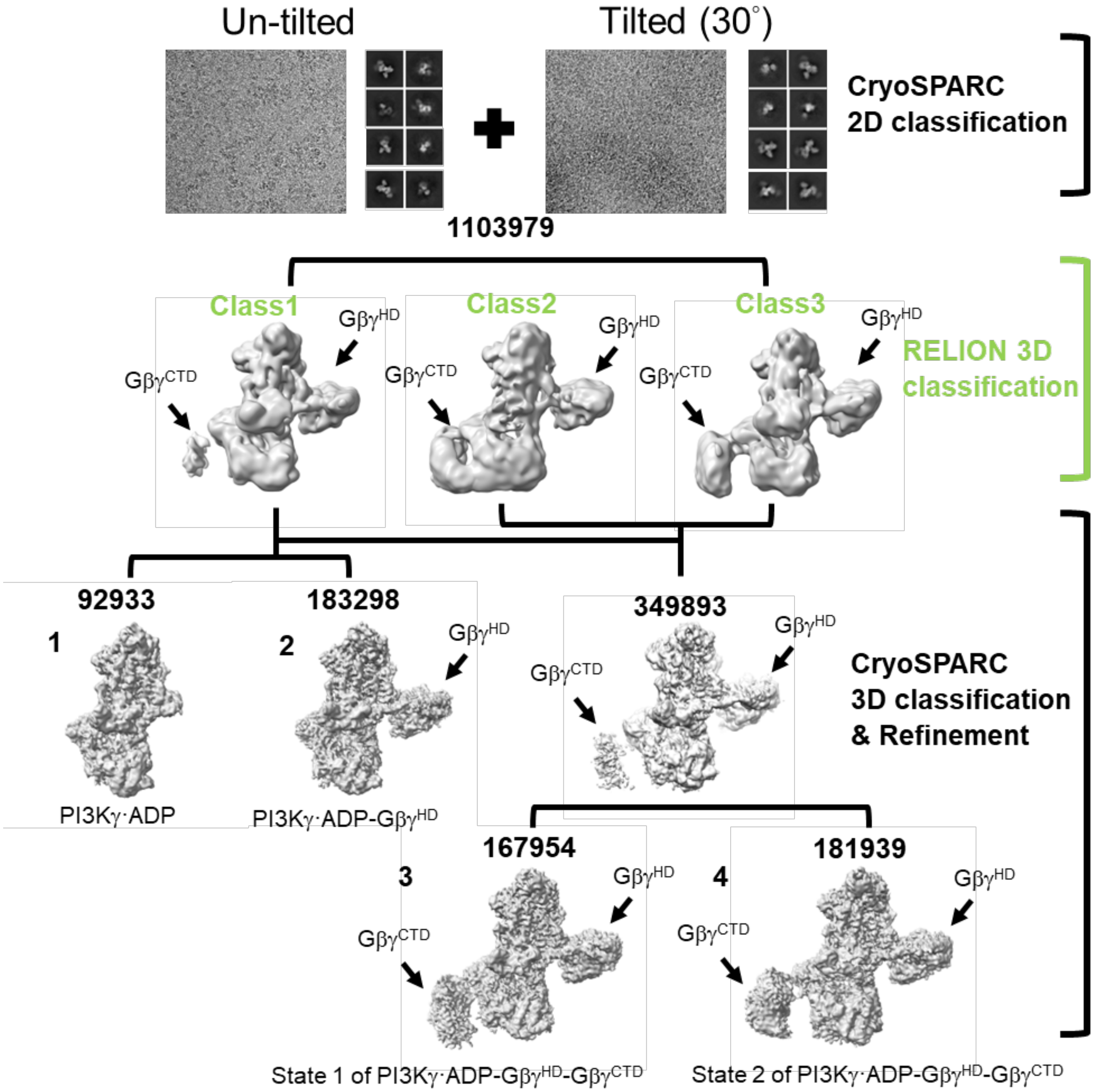


**Extended Data Figure 2. Cryo-EM data processing of PI3Kγ complexed with Gβγ and ADP.** The untilted (3525 frames) and 30° tilted datasets (2943 frames) were collected and combined for data processing in CryoSPARC, resulting in ~1103979 particles for further classification in RELION 3D classification using 3 classes. Particles in Class1 showing weaker $G\beta\gamma^{CTD}$ density were further processed via CryoSPARC 3D classification. Most particles without the $G\beta\gamma^{CTD}$ density classified into two species: PI3Kγ·ADP and PI3Kγ·ADP-$G\beta\gamma^{HD}$. The remaining particles containing the $G\beta\gamma^{CTD}$ were combined with particles from RELION Classes 2 & 3, containing both $G\beta\gamma^{CTD}$ and $G\beta\gamma^{HD}$, for 3D classification in CryoSPARC, resulting in two distinct conformations of PI3Kγ·ADP-$G\beta\gamma^{HD}$-$G\beta\gamma^{CTD}$, State1 and State2, where in $G\beta\gamma^{CTD}$ and the p101 CTD to which it binds exhibiting the largest variation..

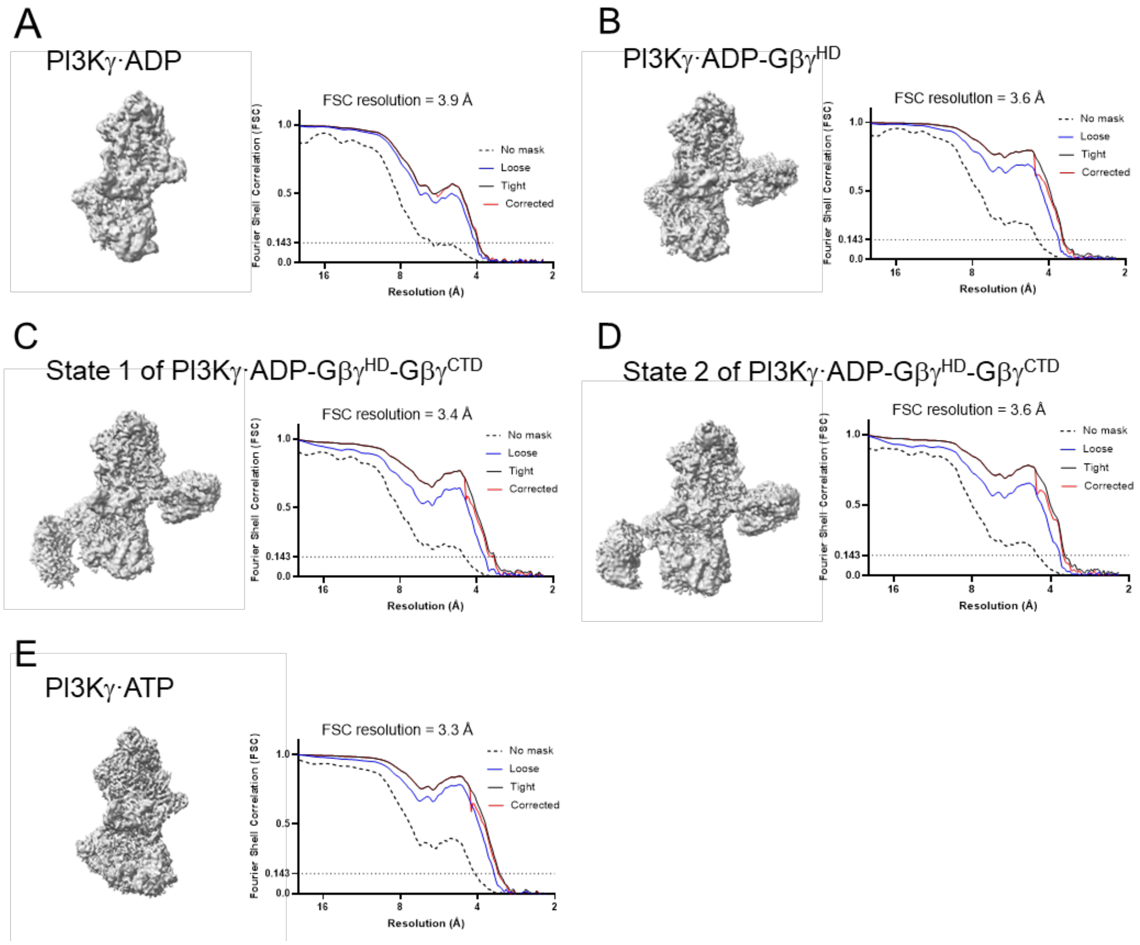


**Extended Data Figure 3. Resolution estimation for cryo-EM reconstructions. (A)** PI3Kγ·ADP: 3.9 Å **(B)** PI3Kγ·ADP-Gβγ$^{HD}$: 3.6 Å **(C)** State 1 of PI3Kγ·ADP-Gβγ$^{HD}$-Gβγ$^{CTD}$: 3.4 Å **(D)** State 2 of PI3Kγ·ADP-Gβγ$^{HD}$-Gβγ$^{CTD}$: 3.6 Å **(E)** PI3Kγ·ATP: 3.3 Å. FSC, Fourier shell correlation.

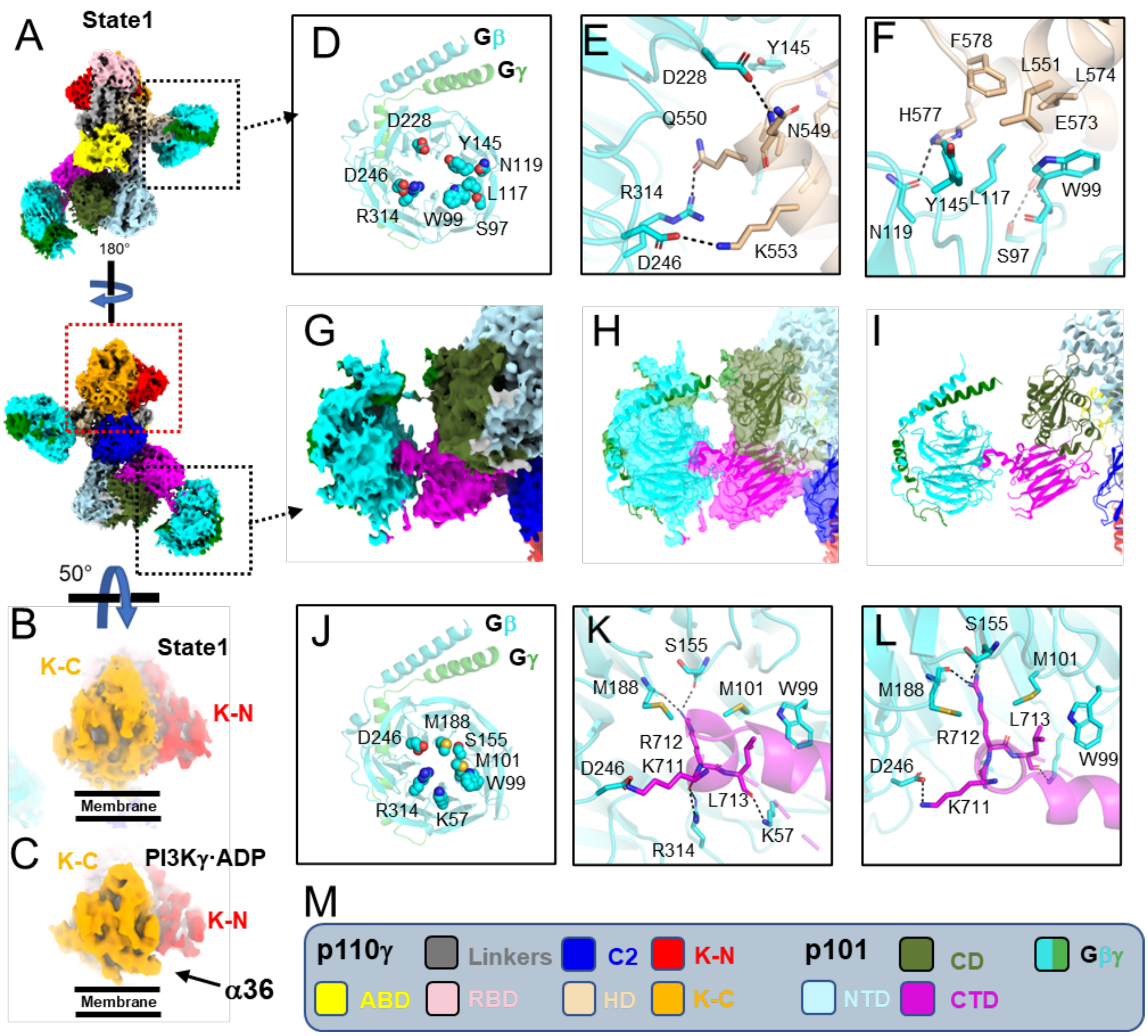


**Extended Data Figure 4. Cryo-EM reconstruction of PI3Kγ·ADP-Gβγ$^{HD}$-Gβγ$^{CTD}$ in State1. (A)** Cryo-EM map of State1 with domain colored and indicated in (M). **(B)** Close-up of the kinase domain in State1 with no observable density for α36. **(C)** Kinase domain in PI3Kγ·ADP, with α36 indicated. **(D)** Gβγ residues involved in the p110γ HD interaction. **(E-F)** Residues in the Gβγ-HD interface. Residues are shown as sticks and colored according to atom type. **(G-I)** Cryo-EM map and model fitting for the p101 CTD-Gβγ$^{CTD}$ interaction. **(J)** Gβγ residues involved in the p110γ CTD interaction. **(K-L)** Residues in the Gβγ-CTD interface. **(M)** Color key for domains in p110γ, p101, and Gβγ.

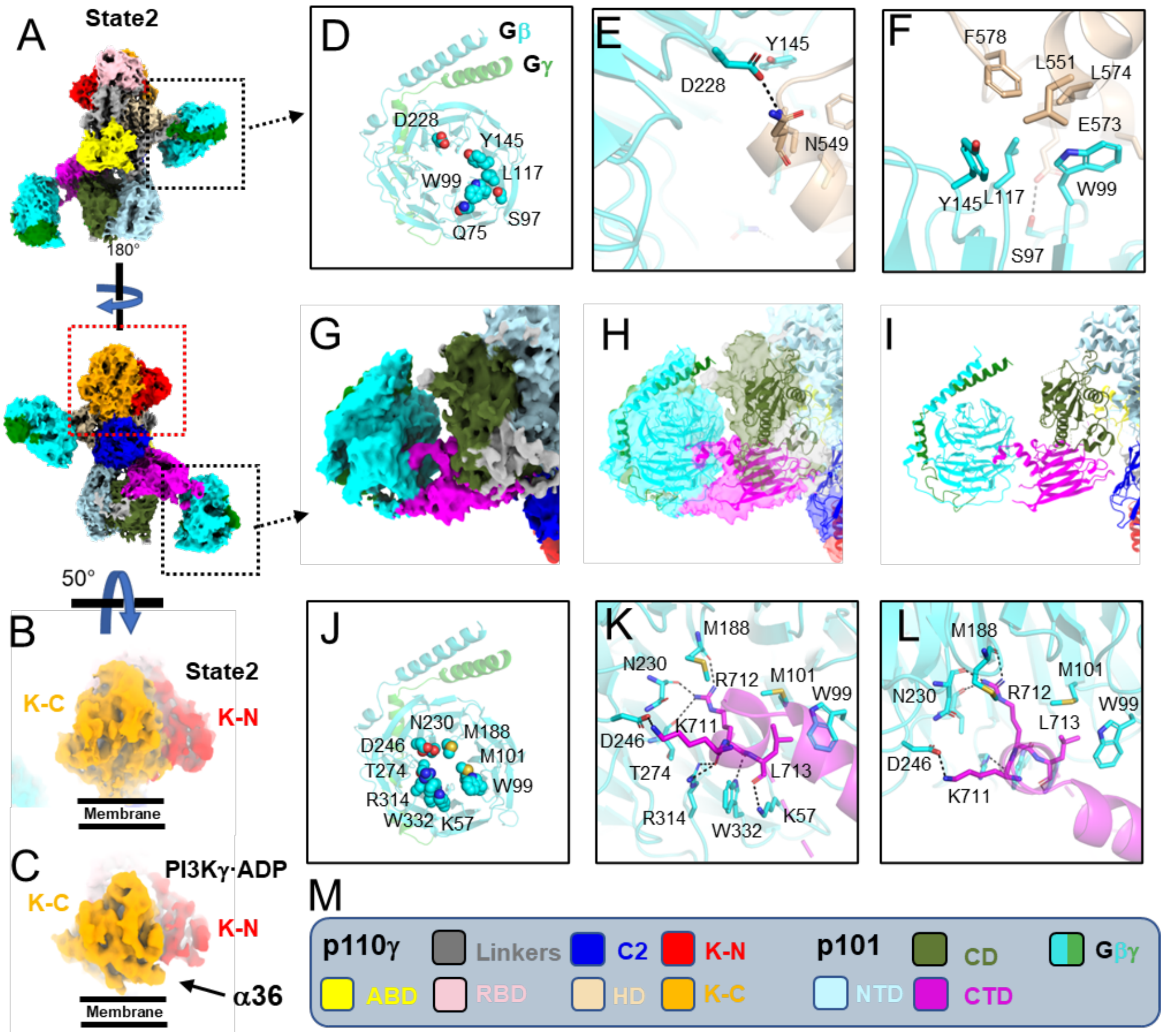


**Extended Data Figure 5. Cryo-EM reconstruction of PI3Kγ·ADP-Gβγ$^{HD}$-Gβγ$^{CTD}$ in State2. (A)** Cryo-EM map of State1 with domain colored and indicated in (M). **(B)** Close-up of the kinase domain in State2 with no observable density for α36. **(C)** Kinase domain in PI3Kγ·ADP, with α36 indicated. **(D)** Gβγ residues involved in the p110γ HD interaction. **(E-F)** Residues in the Gβγ-HD interface. Residues are shown as sticks and colored according to atom type. **(G-I)** Cryo-EM map and model fitting for the p101 CTD-Gβγ$^{CTD}$ interaction. **(J)** Gβγ residues involved in the p110γ CTD interaction. **(K-L)** Residues in the Gβγ-CTD interface. **(M)** Color key for domains in p110γ, p101, and Gβγ.

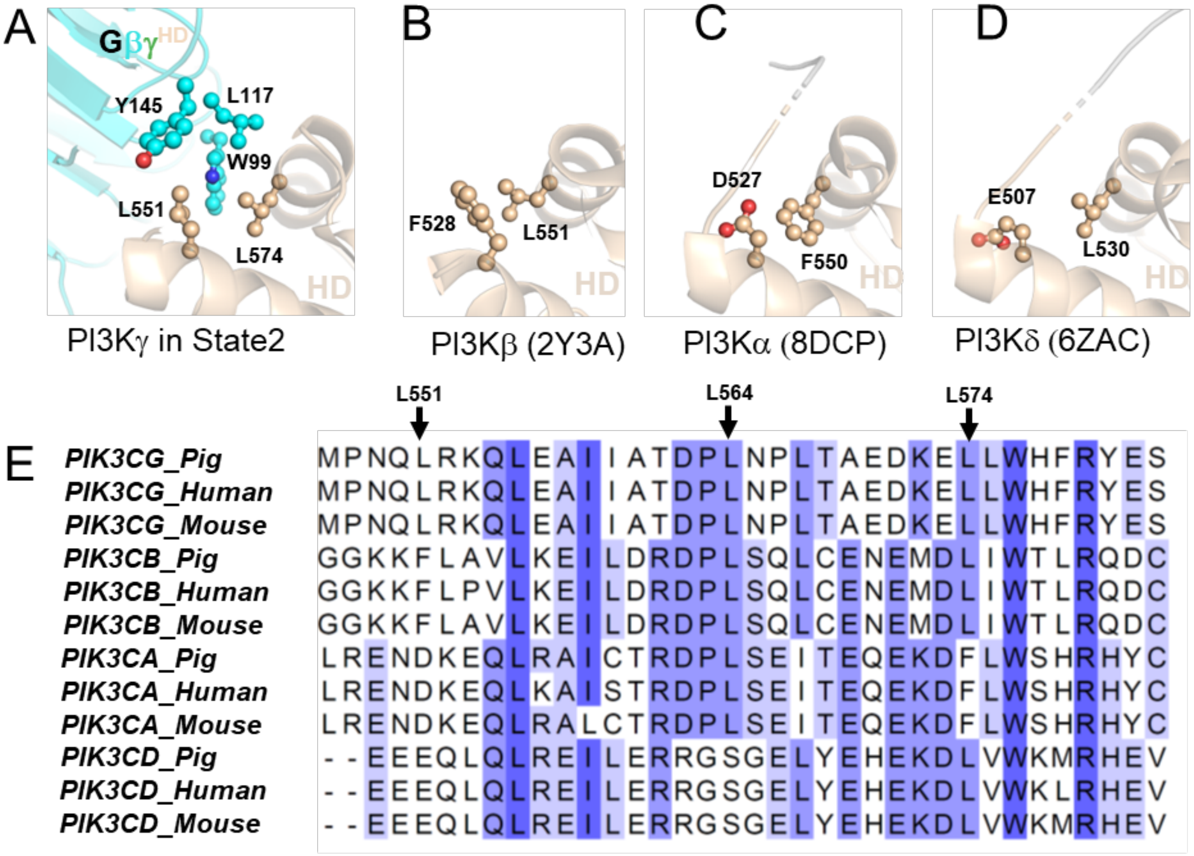


**Extended Data Figure 6. Comparison of PI3K isoforms at the Gβγ$^{HD}$ binding site of PI3Kγ.** Only PI3Kβ retains a hydrophobic signature that may be responsive to Gβγ. **(A)** Hydrophobic interactions between Gβγ$^{HD}$ and p110γ HD. Residues. **(B-D)** Residues of PI3Kβ (PDB entry 2Y3A), PI3Kα (PDB entry 8DCP), and PI3Kδ (PDB entry 6ZAC) at positions corresponding to Leu551 and Leu574 in PI3Kγ are shown as stick models. **(E)** Sequence alignment of p110γ(PIK3CG), β(PIK3CB), α(PIK3CA), and δ(PIK3CD) isoforms (note that human, pig, and mouse sequences are identical within each isoform in this region). The sequence alignment was performed using ClustalW and colored according to BLOSUM-62. Interfacial residues Leu551, Leu564, and Leu574 from pig p110γ are indicated above.

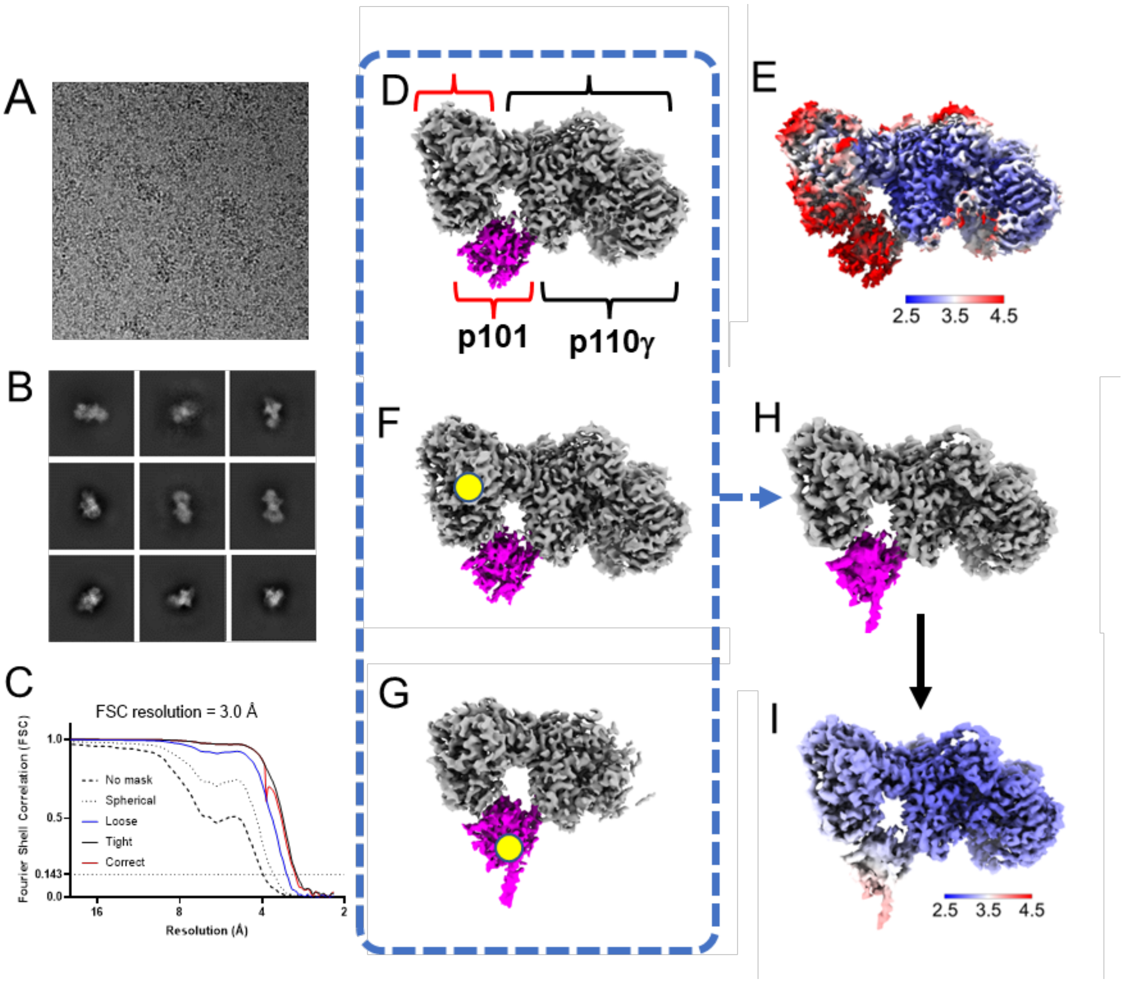


**Extended Data Figure 7. Cryo-EM reconstruction of PI3Kγ. (A)** Representative micrograph. **(B)** Representative 2D class averages (box size = 320 pixels where 1 pixel = 1.08 Å). **(C)** FSC plot, estimating 3.0 Å resolution using FSC at 0.143. **(D)** Cryo-EM reconstruction. In total, 269,357 particles were used for 3D map reconstruction. Map regions belonging to p110γ and p101 are roughly indicated by the black and red brackets. The CTD of p101 is colored magenta. **(E)** Local resolution estimation of the PI3Kγ cryo-EM map from **(D)**. The overall resolution in most regions of p110γ is 2.5-3 Å with well-defined side chains, whereas the overall resolution of p101 is greater than 3.5 Å. **(F)** Local refinement of PI3Kγ focused on the p101 NTD. The yellow circle indicates the fulcrum point used for local refinement. **(G)** Local refinement of PI3Kγ focused on the p101 C-terminal domain. The yellow circle indicates the fulcrum point used for local refinement. **(H)** Composite map generated from the three maps shown in panels **(D)**, **(F),** and **(G)**. **(I)** Local resolution estimation of the composite PI3Kγ cryo-EM map from **(H)**. The overall resolution in most regions is 2.5-3 Å, whereas the p101 C-terminal domain has an improved resolution at 3-4 Å.

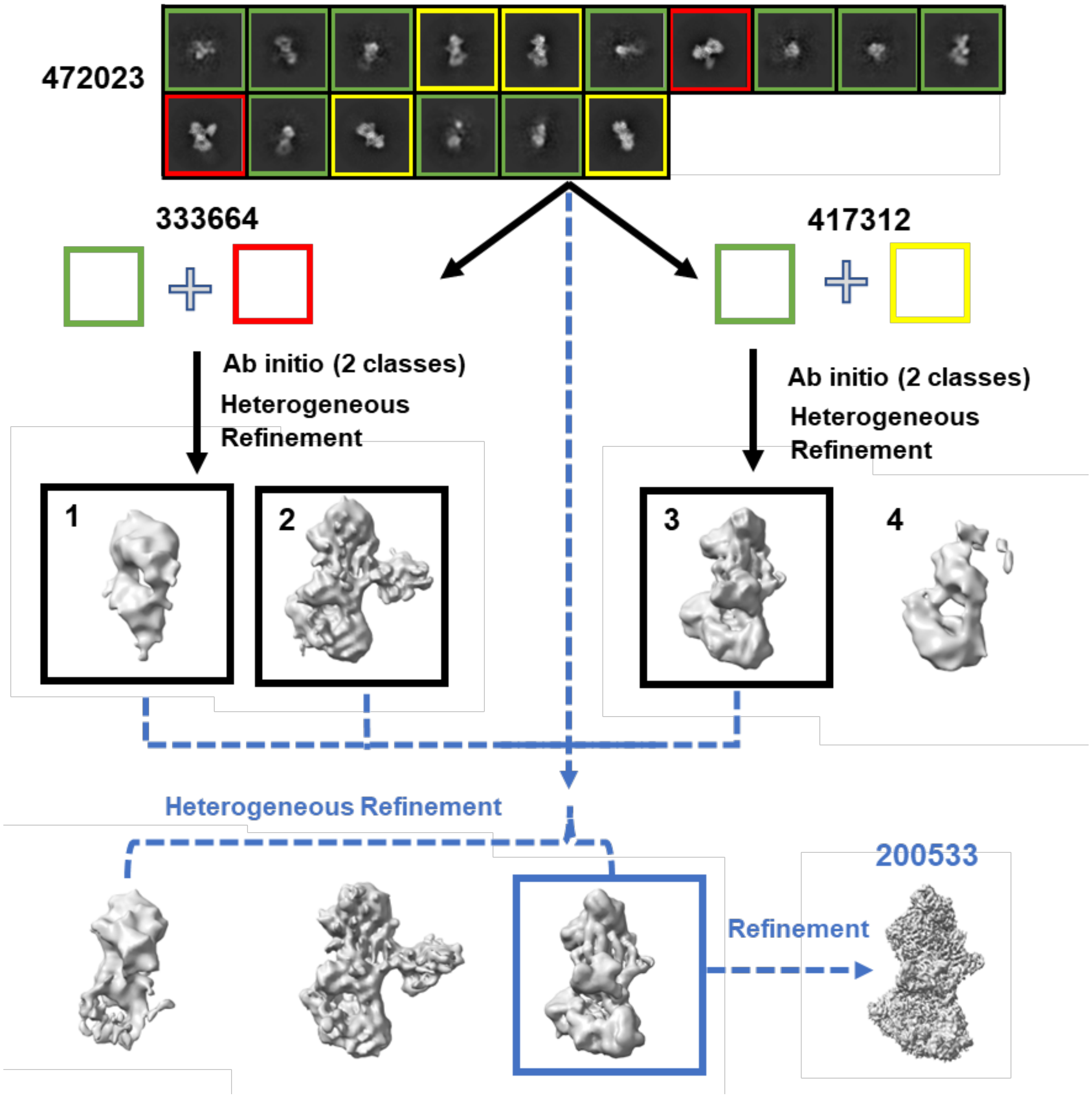


**Extended Data Figure 8. Cryo-EM data processing of PI3Kγ·ATP derived from an un-tilted dataset of PI3Kγ supplemented with Gβγ and ATP.** We collected 3936 frames for the PI3Kγ sample supplemented with Gβγ and ATP and processed the data in CryoSPARC, resulting in 472,023 particles after several rounds of 2D classification. The 2D classes showing distinct Gβγ density are indicated by red boxes. The 2D classes showing no Gβγ density are indicated by yellow boxes. The classes in green boxes are undetermined. We generated 2 *ab initio* maps using the particles from the "red" and "green" 2D classes, followed by heterogeneous refinement in CryoSPARC, resulting in a "junk particle" and "Gβγ-bound PI3Kγ" cryo-EM maps. In parallel, we processed the particles from the "yellow" and "green" 2D classes using the same strategy, resulting in a "PI3Kγ alone" and another "junk particle" cryo-EM map. We then used the 3 maps (1: junk-particle, 2:PI3Kγ-Gβγ, and 3:PI3Kγ alone) for sorting all particles using heterogeneous refinement and did homogeneous and local refinements for the "PI3Kγ alone" data, resulting in a 3.3 Å cryo-EM reconstruction of PI3Kγ·ATP (**Extended Data Fig. 3E**).

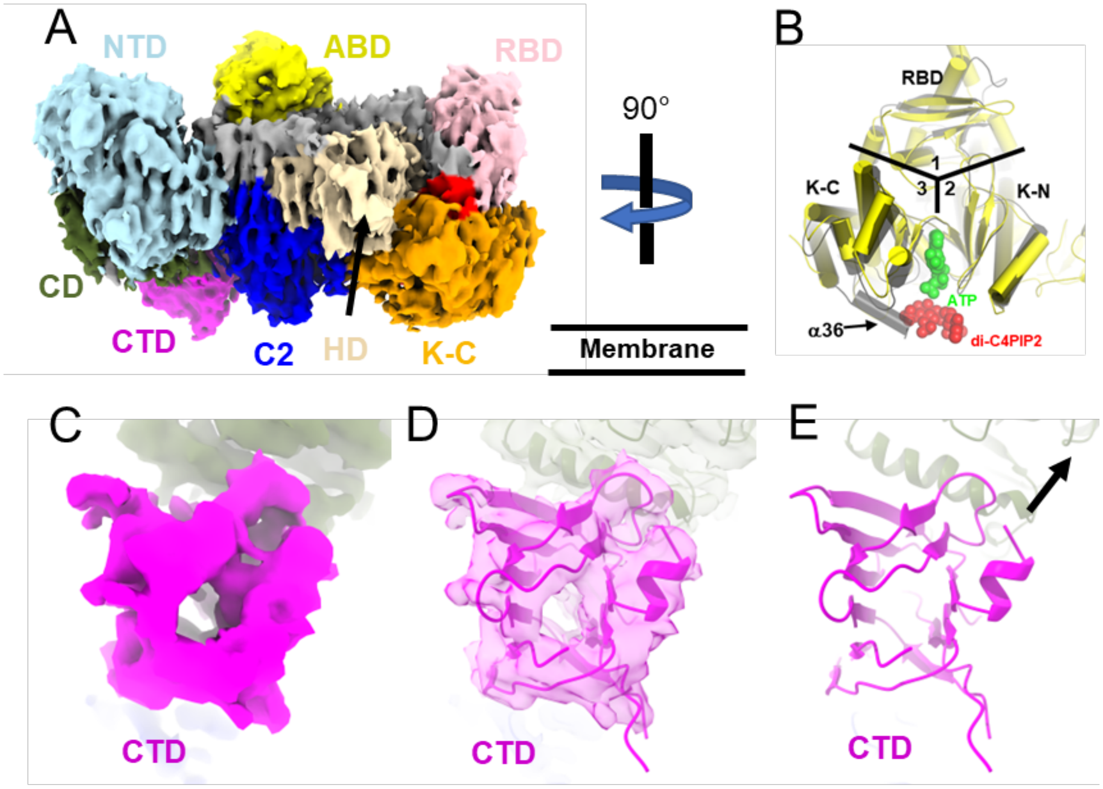


**Extended Data Figure 9. Cryo-EM reconstruction of PI3Kγ·ATP. (A)** Cryo-EM map of PI3Kγ·ATP. Domains are colored as in **Fig 1A**. **(B)** Superimposition of PI3Kα·diC4-PIP2 (PDB entry 4OVV, in yellow) with PI3Kγ·ATP (in black) shows a steric clash between the $PIP_2$ substrate and α36. Molecules, diC4-$PIP_2$ and ATP in PI3Kγ·ATP are shown as spheres colored green and red, respectively. The RBD, K-N, and K-C domains are located in areas 1, 2, and 3, respectively. **(C-E)** Close up view of the p101 CTD. In (E), α18 (residues 695-701) is indicated by a black arrow in the direction from its N- to C-terminal ends.

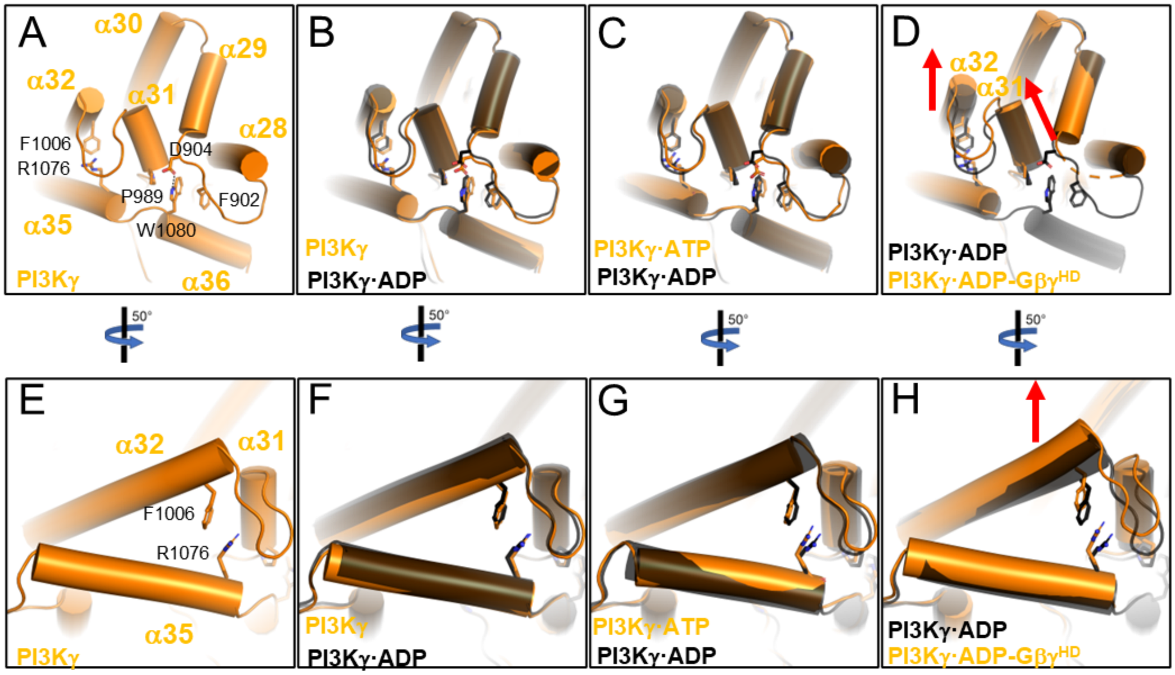


**Extended Data Figure 10. Comparison of the kinase domain C-lobe in PI3Kγ, PI3Kγ·ADP, PI3Kγ·ATP, and PI3Kγ·ADP-Gβγ$^{HD}$ (A-D)** The kinase domain C-lobe in various PI3Kγ structures. In **(A)**, the tryptophan lock, W1080, and the interacting residues F902, D904, and P989 are shown with stick side chains. **(E-H)** A close up images of α32 and α35 and residues Phe1006 and Arg1076, which form a π-stacking interaction in less activated states.

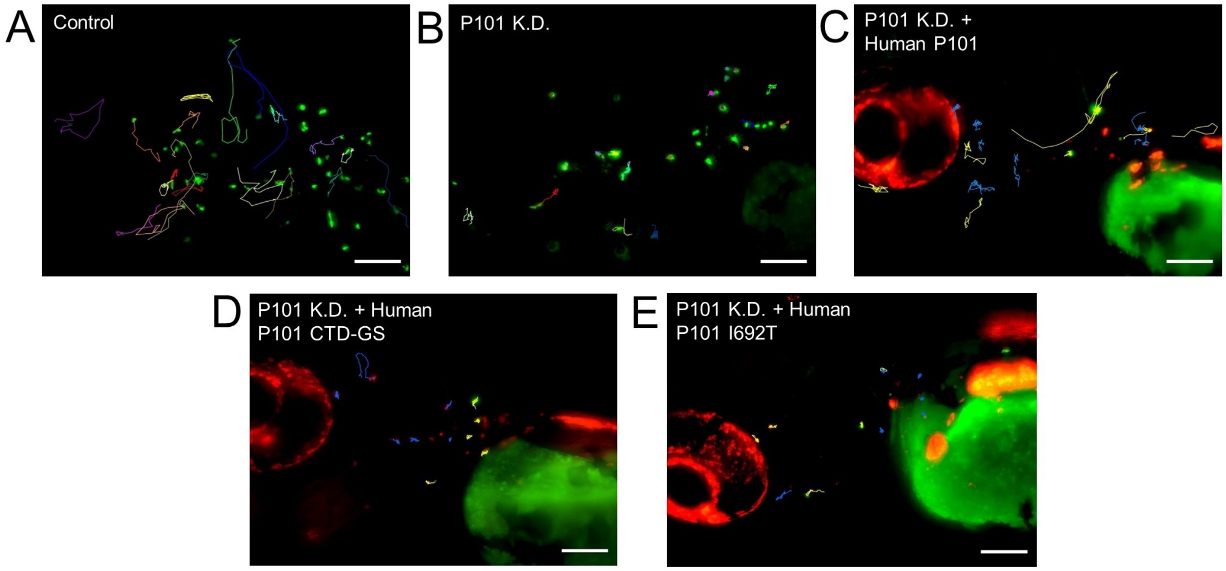


**Extended Data Figure 11. Neutrophil migration tracking. (A-E)** Neutrophil migration imaging in a zebrafish embryo 3 days post-fertilization. **(A-B)** Representative images of neutrophil tracks under control (PBS) or P101 knockdown (MO) background. **(C-E)** Representative images of neutrophils expressing human P101 (**C)**, mutant human P101 CTD-GS **(D)**, or mutant human P101 I690T **(E)** indicated by yellow tracks under P101 knockdown background (blue tracks). Scale bar (white): 100 µm.

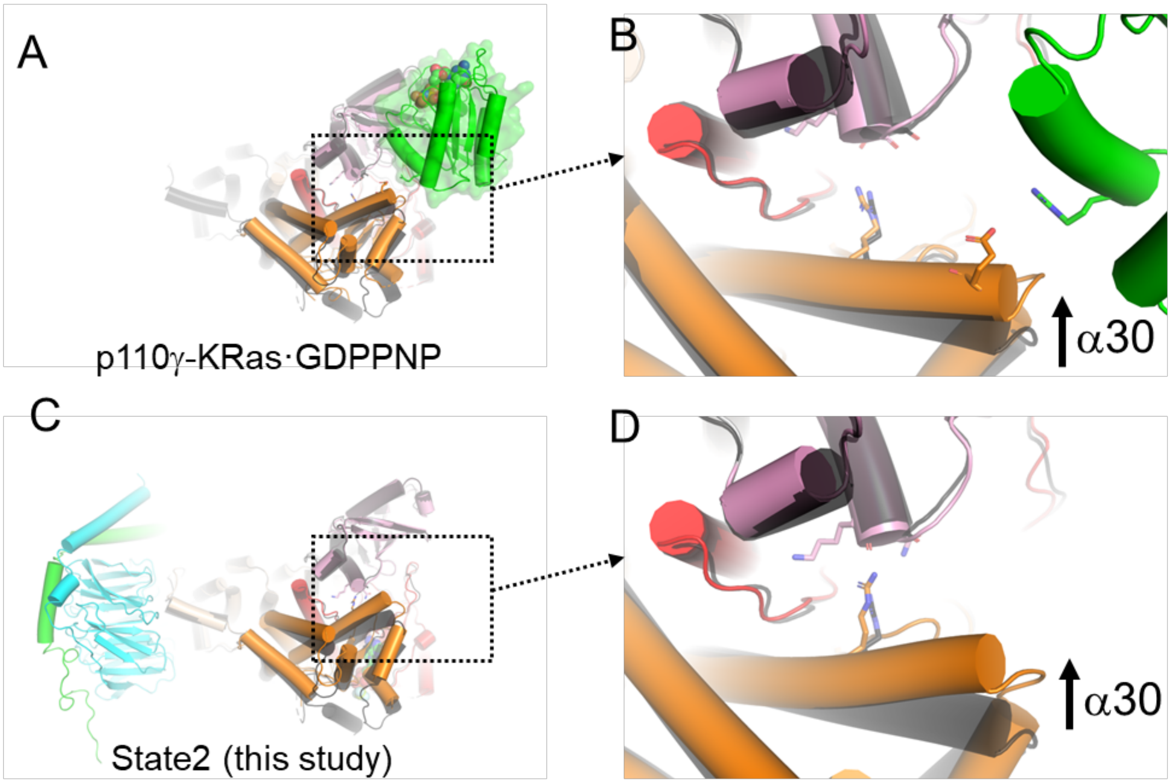


**Extended Data Figure 12. Gβγ- and KRas-mediated conformational changes in PI3Kγ are similar. (A)** Superposition of PI3Kγ (black) with the crystal structure of the p110γ subunit complexed with activated K-Ras (p110γ–K-Ras·GDPPNP, PDB entry 1HE8). Activated K-Ras bound with GDPPNP is shown in green cartoon representation surrounded by a green transparent surface. **(B)** A close-up view of the kinase domain N-lobe. A shift of α30 is indicated. **(C)** Superposition of PI3Kγ (black) with State 2 of PI3Kγ·ADP–Gβγ$^{HD}$–Gβγ$^{CTD}$. **(D)** A close-up view in the kinase domain N-lobe. A shift of α30 is indicated.

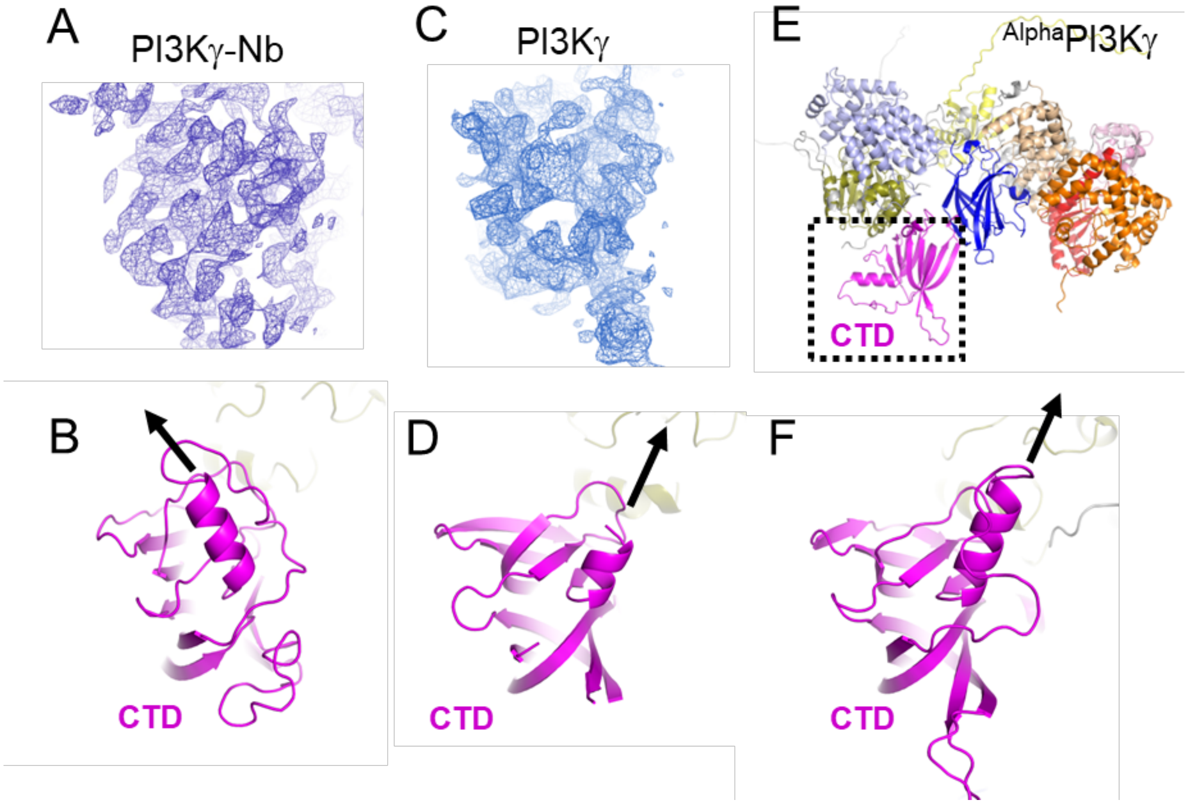


**Extended Data Figure 13. Comparison of maps and models of PI3Kγ, PI3Kγ-Nb, and Alphafold predicted PI3Kγ ($^{Alpha}$PI3Kγ) around the p101 CTD. (A)** Cryo-EM map of $^{HP}$PI3Kγ-Nb (PDB entry 7MEZ) p101 CTD**. (B)** Atomic model thereof. **(C)** Cryo-EM map of PI3Kγ (PDB entry 8SO9) p101 CTD**. (D)** Atomic model thereof. **(E)** Cartoon representation of $^{Alpha}$PI3Kγ with domain colored as described in Fig 1A. **(F)** A zoom-in view of the p101 C-terminal domain in the same orientation as that in **(A)-(D). In (B)**, **(D),** and **(F)**, the helix orientation of α18 of PI3Kγ (residues 695-701) is indicated by black arrows pointing from their N- to C- terminal ends.

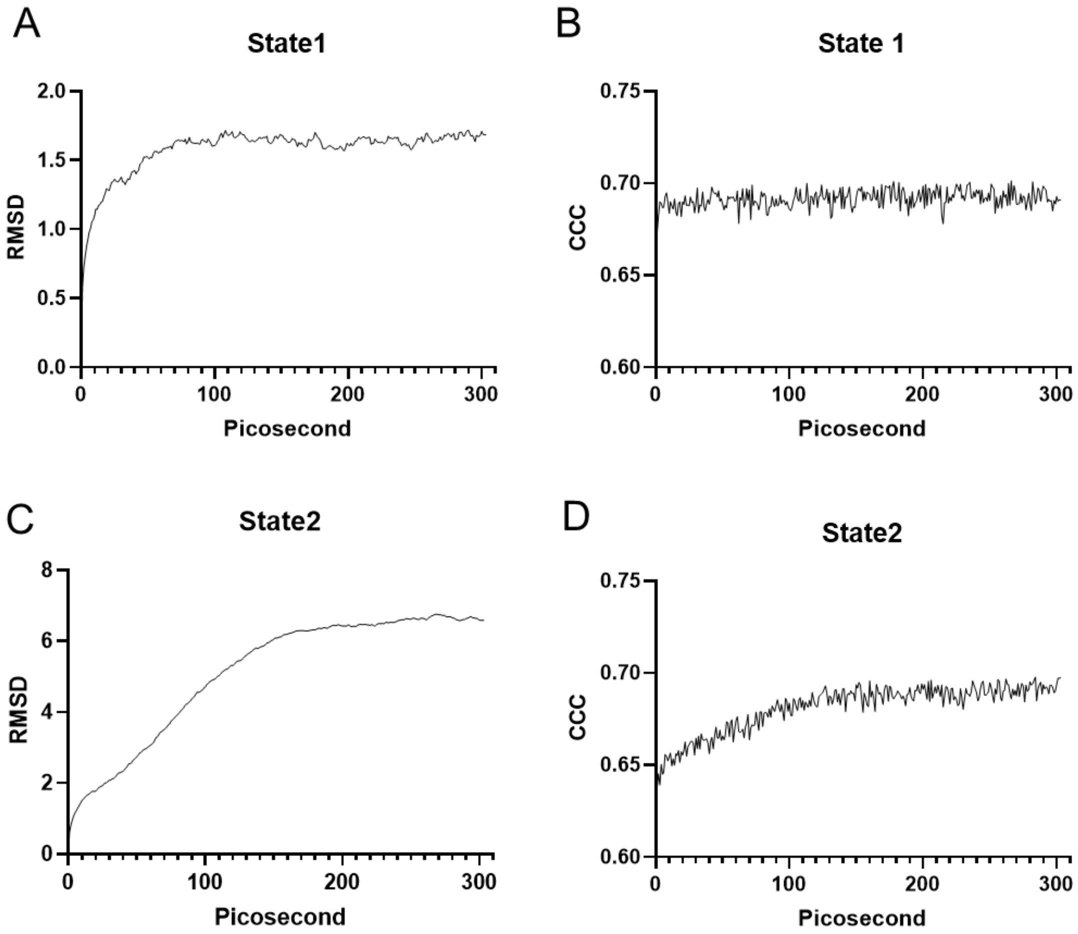


**Extended Data Figure 14. Molecular dynamics flexible fitting for the two conformational states of PI3K$\gamma$·ADP–G$\beta\gamma^{HD}$–G$\beta\gamma^{CTD}$. (A-B)** Calculated C$\alpha$ RMSD and the cross-correlation coefficient (CCC) of the simulated structures to the starting model for State 1. **(C-D)** Calculated C$\alpha$ RMSD and the CCC of the simulated structures to the starting model for State 2. See Methods.

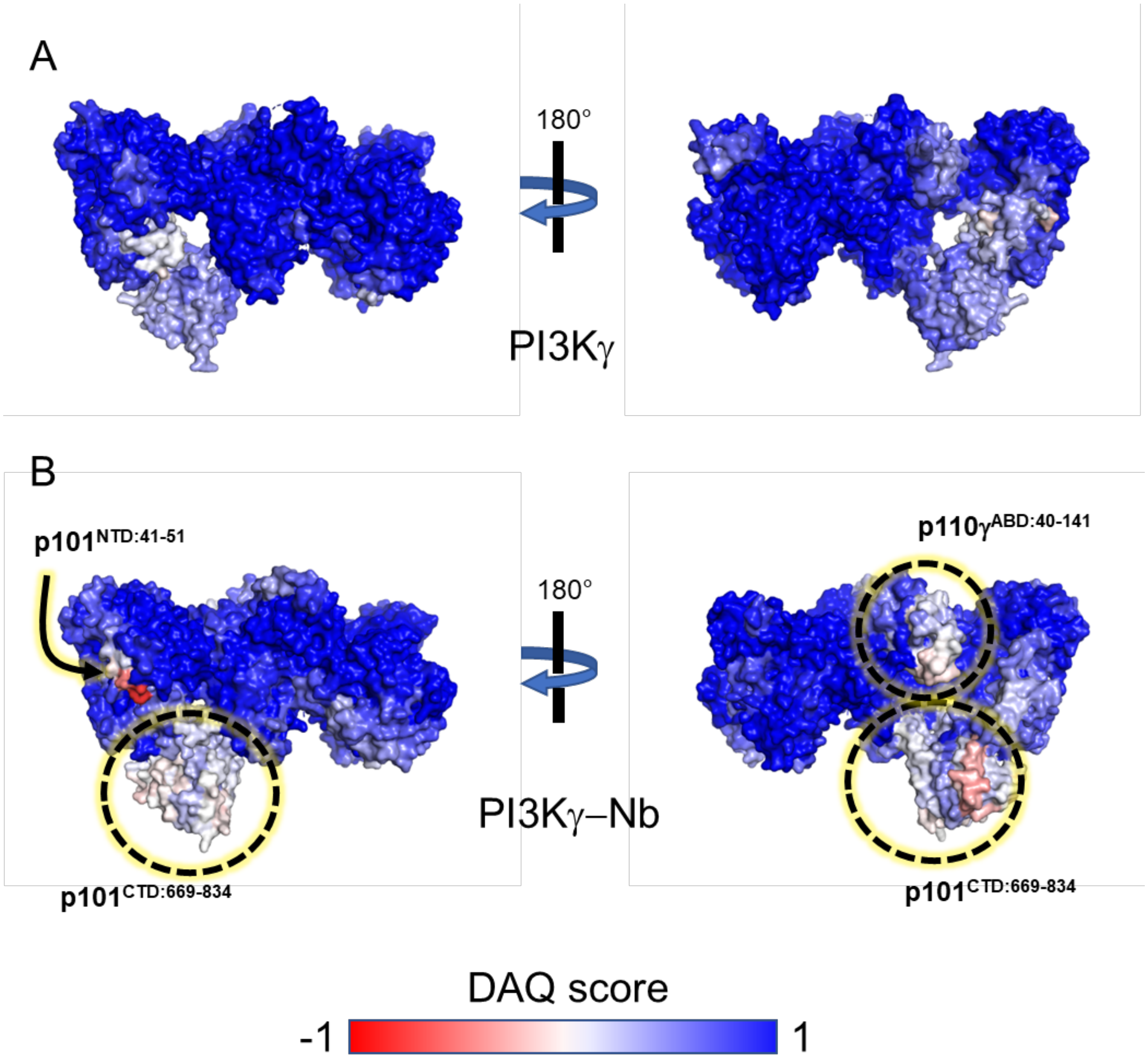


**Extended Data Figure 15. Residue-wise local quality estimation (DAQ) of PI3Kγ and PI3Kγ–Nb. (A)** The DAQ evaluation of the PI3Kγ model (PDB entry 8SO9) to the corresponding cryo-EM map in this study. **(B)** DAQ evaluation of the PI3Kγ-Nb model (PDB entry 7MEZ) to its corresponding cryo-EM map (EMDB entry EMD-23808). Regions with poor model/map agreement (DAQ < 0) are labeled and indicated by an arrow and dashed circles.

### Extended Data Table 1 Structure Statistics

| | PI3Kγ | PI3Kγ·ATP | PI3Kγ·ADP | PI3Kγ·ADP-$G\beta\gamma^{HD}$ | State1 of PI3Kγ·ADP-$G\beta\gamma^{HD}$-$G\beta\gamma^{CTD}$ | State2 of PI3Kγ·ADP-$G\beta\gamma^{HD}$-$G\beta\gamma^{CTD}$ |
|---|---|---|---|---|---|---|
| **PDB** | 8SO9 | 8SOA | 8SOB | 8SOC | 8SOD | 8SOE |
| **EMDB** | EMD-40650 | EMD-40651 | EMD-40652 | EMD-40653 | EMD-40654 | EMD-40655 |
| **Data collection and processing** | | | | | | |
| **Magnification** | 81000 | 81000 | 81000 | 81000 | 81000 | 81000 |
| **Voltage (kV)** | 300 | 300 | 300 | 300 | 300 | 300 |
| **Electron exposure (e–/Å$^2$)** | 56 | | | | | |
| **Defocus range (μm)** | -0.5~-2.2 mm | -0.5~-2.2 mm | -0.5~-2.2 mm | -0.5~-2.2 mm | -0.5~-2.2 mm | -0.5~-2.2 mm |
| **Pixel size (Å)** | 0.54 | 0.54 | 0.54 | 0.54 | 0.54 | 0.54 |
| **Symmetry imposed** | C1 | C1 | C1 | C1 | C1 | C1 |
| **Initial particle images (no.)** | | | | | | |
| **Final particle images (no.)** | 269,357 | 200,533 | 92,933 | 183,298 | 167,954 | 181,939 |
| **Map resolution (Å)**<br>**(FSC threshold=0.143)** | 3.0 | 3.3 | 3.9 | 3.6 | 3.4 | 3.6 |
| **Refinement** | | | | | | |
| **Map sharpening B factor (Å$^2$)** | 122.0 | 118.2 | 145.2 | 135.1 | 134.1 | 136.5 |
| **Model composition**<br>**Non-hydrogen atoms**<br>**Protein residues** | 12250 / 1523 | 11975 / 1481 | 12163 / 1507 | 14872 / 1860 | 18050 / 2271 | 17789 / 2238 |
| **B factors (Å$^2$)**<br>**Protein**<br>**Ligand** | 172.6 / --- | 84.1 / 67.74 | 148.5 / 141.0 | 152.1 / 149.7 | 163.5 / 137.5 | 194.7 / 160.0 |
| **R.m.s. deviations**<br>**Bond lengths (Å)**<br>**Bond angles (°)** | 0.004 / 0.6 | 0.004 / 0.571 | 0.005 / 0.587 | 0.004 / 0.608 | 0.004 / 0.701 | 0.003/ 0.670 |
| **Validation**<br>**MolProbity score/**<br>**Clashscore/**<br>**Poor rotamers (%)** | 1.77 / 2.52 / 2.79 | 1.73 / 3.37 / 2.94 | 2.07 / 5.37 / 2.89 | 1.90 / 4.91 / 2.26 | 2.32 / 7.69 / 3.33 | 2.13 / 5.24 / 3.53 |
| **Ramachandran plot**<br>**Favored (%)/**<br>**Allowed (%)/**<br>**Disallowed (%)** | 94.1 / 5.7 / 0.2 | 96.33 / 3.6 / 0.07 | 93.29 / 6.64 / 0.07 | 94.44 / 5.45 / 0.11 | 91.62 / 8.34 / 0.04 | 93.28 / 6.68 / 0.04 |

#### Extended Data Table 2. Primers for mutagenesis of *Sus scrofa* PI3Kγ

| Primer | Forward (F) or Reverse (R) | Sequence |
|---|---|---|
| p110γ | F | CCCGGAGAATTCATGCATCACCATCACCATCACGAGCTGGAGAACTATGAACAGC |
| p110γ | R | CCCAGGTCTAGATTATGCGGAATGCTTCTCCCCTTGTTTGATGCC |
| p101 | F | AGCCATGGTGCTAGCATGCAGCCAGGGGCCACGACGTGCACGGAGGACCGC |
| p101 | R | TCTCCCGGTACCCTACTTATCGTCGTCATCCTTGTAATCAGAGCC |
| p110γ $^{L551A}$ | F | GCCCAATCAGGCACGGAAGCAACTGG |
| p110γ $^{L551A}$ | R | CCAGTTGCTTCCGTGCCTGATTGGGC |
| p110γ $^{L564S}$ | F | CACGGATCCGAGCAACCCACTCAC |
| p110γ $^{L564S}$ | R | GTGAGTGGGTTGCTCGGATCCGTG |
| p101 $^{I689T}$ | F | GGAGATCTTCACCCACTCCCTGG |
| p101 $^{I689T}$ | R | CCAGGGAGTGGGTGAAGATCTCC |
| p101 $^{CTD-GS}$ | F | CAAGGCTTCGGGTCCTGGCAGCGGCAGCCTGGGCATCGATGGTGACCGGGAGGCCGTCC |
| p101 $^{CTD-GS}$ | R | GGACGGCCTCCCGGTCACCATCGATGCCCAGGCTGCCGCTGCCAGGACCCGAAGCCTTG |
| p101 $^{CTD-GSS}$ | F | CAAGGCTTCGGGTCCTGGCAGCGGCAGCAGTGGCATCGATGGTGACCGGGAGGCCGTCC |
| p101 $^{CTD-GSS}$ | R | GGACGGCCTCCCGGTCACCATCGATGCCACTGCTGCCGCTGCCAGGACCCGAAGCCTTG |

#### Extended Data Table 3. Specific activity of PI3Kγ variants

| Complex Name | Catalytic Subunit | Regulatory Subunit | Basal Specific Activity (nmol ADP*μmol enzyme$^{-1}$*min$^{-1}$) | Maximum Specific Activity (Gβγ) (nmol ADP*μmol enzyme$^{-1}$*min$^{-1}$) | Fold Activation | n |
|---|---|---|---|---|---|---|
| WT | p110γ | p101 | 900± 400 | 9000± 400 | 10 | 8 |
| L551A | L551A | p101 | 850± 100 | 2100± 100 | 2.4 | 14 |
| L564S | L564S | p101 | 850± 400 | 2200± 200 | 2.6 | 8 |
| I689T | p110γ | I689T | 730± 200 | 2100± 300 | 2.9 | 10 |
| CTD-GS | p110γ | 711GS | 970± 200 | 4400± 200 | 4.5 | 8 |
| CTD-GSS | p110γ | 711GSS | 1200± 200 | 4400± 400 | 3.6 | 8 |

* Enzymatic reactions were conducted at room temperature for 40 minutes, followed by measuring ADP production with ADP-glo. The values of basal and maximum specific activity were derived from curve fitting as shown in **Fig 5C-F** (see methods).

**Extended Data Movie 1. Tracking movies of migrating neutrophils in the head mesenchyme with p101 knockdown.** The video shows the motility of neutrophils in 3 dpf (days post fertilization)*Tg(lyzC:RFP)* zebrafish larvae injected with buffer or p101 morpholino. Videos were recorded for 30 min at 1 min intervals.

Representative videos from n = 3 independent experiments with four fish in each group are shown. Scale bar: 100 μm.

**Extended Data Movie 2. Tracking movies of migrating neutrophils in the head mesenchyme re-expressing p101 wild type or mutations in the p101 knockdown background.** The videos show the motility of neutrophils in 3 dpf *Tg(lyzC:RFP*) zebrafish larvae injected with buffer or p101 morpholino, then injected with Tol2-lyzC-human p101, mutant human p101 I692T, or p101 CTD-GS plasmids for transient re-expression. Videos were recorded for 30 min with 1 min intervals. Yellow tracks indicate neutrophils with transient p101 expression. Blue tracks indicate neutrophils with p101 knockdown in the same fish. Representative videos from n = 3 independent experiments with four fish in each group are shown. Scale bar: 100 μm.
